# SETD2 promotes PAF1C interactions with the elongating RNA Pol II and is required for neuronal differentiation

**DOI:** 10.1101/2024.10.23.619798

**Authors:** Christina Ambrosi, Ramon Pfaendler, Stefan Butz, Davide Recchia, Xue Bao, Nina Schmolka, Tuncay Baubec

**Affiliations:** Department of Molecular Mechanisms of Disease, University of Zurich, Zurich, Switzerland; Life Science Zurich Graduate School, University of Zurich and ETH Zurich, Zurich, Switzerland; Genome Biology and Epigenetics, Institute of Biodynamics and Biocomplexity, Department of Biology, Utrecht University, Utrecht, The Netherlands

**Author notes:** Institute of Molecular Systems Biology, ETH Zurich, Zurich, Switzerland. Institute of Experimental Immunology, University of Zurich, Zurich, Switzerland (Nina Schmolka).

## Abstract

Chromatin modifications are relevant for mammalian development and their aberrant deposition is associated with human disease. While the mechanisms that deposit and remove these modifications have been largely elucidated, their role in regulating gene activity during cellular differentiation have yet to be completely understood. By differentiating a panel of mouse embryonic stem cells lacking major chromatin regulators towards neuronal cells, we identified their requirement at different stages of cellular differentiation. We show that the H3K36me3 methyltransferase SETD2 is important for the establishment of neuronal gene expression during late stages of differentiation, but is dispensable once the cells have fully differentiated. This function is largely independent of the histone methyltransferase activity. By measuring the protein interaction network of elongating RNA Pol II, we identify a novel role for SETD2 in mediating interactions between the PAF1 complex and the elongating RNA Pol II, which is required to ensure optimal transcriptional processivity of neuronal genes.

## Introduction

Chromatin modifications play important roles in orchestrating site-specific processes in a genomic context. Mutations in the enzymes that deposit these modifications and aberrant distribution of chromatin marks result in impaired development and are associated with numerous human diseases ^1,2^. Thus, chemical marks on DNA and histones contribute to the establishment and maintenance of correct gene expression patterns and are essential for cellular identity and function. In the last decades, the genomic distribution of most chromatin marks has been characterized in various cell types and organisms, allowing us to better understand their potential involvement in regulating transcriptional programs ^3,4^.

Within the large panel of chemical modifications of chromatin, methylation marks on the histone H3 tails at lysine residues K9 or K27, and/or methylation of cytosines on the DNA are frequently associated with transcriptional inactivity of gene promoters and repetitive elements ^5,6^. Besides these repressive modifications, tri-methylation of histone H3 at residue K36 (H3K36me3) is deposited by the methyltransferase SETD2 through interactions with the elongating RNA Polymerase II ^7^, resulting in the bookmarking of transcribed genes by the H3K36me3 mark. The role of this bookmarking is not fully understood in mammals and several regulatory functions have been reported, including prevention of spurious transcription initiation from gene bodies ^8^, DNA damage response ^9^, splicing regulation ^10^, and chromatin cross-talk ^11,12^. Loss of H3K36me3, either through mutations in *Setd2* or histone H3 K36M mutations, as observed in clear cell renal cell carcinoma ^13,14^ and chondrosarcoma ^15,16^, respectively, hint to a potential role of this modification in preventing transformation and undifferentiated growth – highlighting its relevance for cellular function.

Interestingly, despite the functional requirement for SETD2 to catalyse the deposition of H3K36me3, ablation of *Setd2* in embryonic stems cells results in limited changes in gene expression and cellular identity, which is in stark contrast to the phenotypes observed in *Setd2*-KO mouse models ^12,17,18^. Similar outcomes were reported for other important chromatin modifying enzymes, including H3K27me3 or DNA methylation writers, where stem cells could survive in absence of these marks, while their differentiation was impaired ^19–21^. This apparent cellular context-dependent role of chromatin modifications remains to be fully elucidated and requires systematic comparisons to test the contribution of individual chromatin modifications towards the establishment and maintenance of cell type-specific gene expression. Furthermore, recent studies highlight a role for histone-modifying enzymes that expands beyond their catalytic activities, where lack of catalytic activity was dispensable, questioning the essentiality of some of the histone modifications ^22^.

In this study we compare the neuronal differentiation potential of isogenic mESC lines lacking the major chromatin regulators EED, DNMT1, DNMT3A, DNMT3B, and SETD2 to test their requirements for exit from pluripotency, lineage commitment, and terminal differentiation. We identify SETD2 to be important for the establishment of neuronal gene expression patterns, but dispensable once these are established. We show that this function does not require the catalytic activity of SETD2, suggesting a non-catalytic role for this protein. By employing ChromID ^23^ to identify the proteins associated with the elongating RNA Pol II in absence and presence of SETD2, we reveal a reduced association of the PAF1 Complex with the elongating polymerase. These results suggest a non-catalytic role of SETD2 in mediating interactions between RNA Pol II and PAF1C for promoting efficient transcription of long neuronal genes.

## Results

### Stage-specific requirement for chromatin regulators indicates a role for SETD2 in neuronal differentiation

To compare the contribution of DNA methylation, H3K27me3, and H3K36me3 to gene expression during cellular differentiation, we compiled a set of CRISPR-generated knock-out mouse embryonic stem cell (mESC) lines including *Dnmt1,3a,3b* triple-KO (“TKO”) ^24^, *Eed*-KO ^23^, and *Setd2*-KO ^12^, all created in the same isogenic background (Figure 1a). In addition, to test the combinatorial contribution of DNA methylation and H3K36me3, we ablated *Setd2* in the TKO background (quadruple-KO, “QKO”) (Supplementary Figure 1a-d). All cell lines retained pluripotency and were able to proliferate in serum + LIF conditions without apparent changes in morphology, proliferation capacity, and lineage marker expression (Supplementary Figure 2a-e). Using these cell lines, we wanted to compare how the absence of these chromatin modifications influence exit from pluripotency and lineage commitment. Towards this, we differentiated the mESCs to neural progenitors and subsequently to maturing glutamatergic neurons *in vitro* ^25^ (Figure 1b). As expected from previous studies ^17,19–21^ differentiation in cell lines lacking DNA methylation and H3K27me3 was severely compromised since *Eed*-KO cells did not survive exit from pluripotency (CAd4) and the *Dnmt-*TKO cells aborted differentiation at the neural progenitor stage (NPC) (Figure 1b and Supplementary Figure 3a-b).

**Figure 1.**
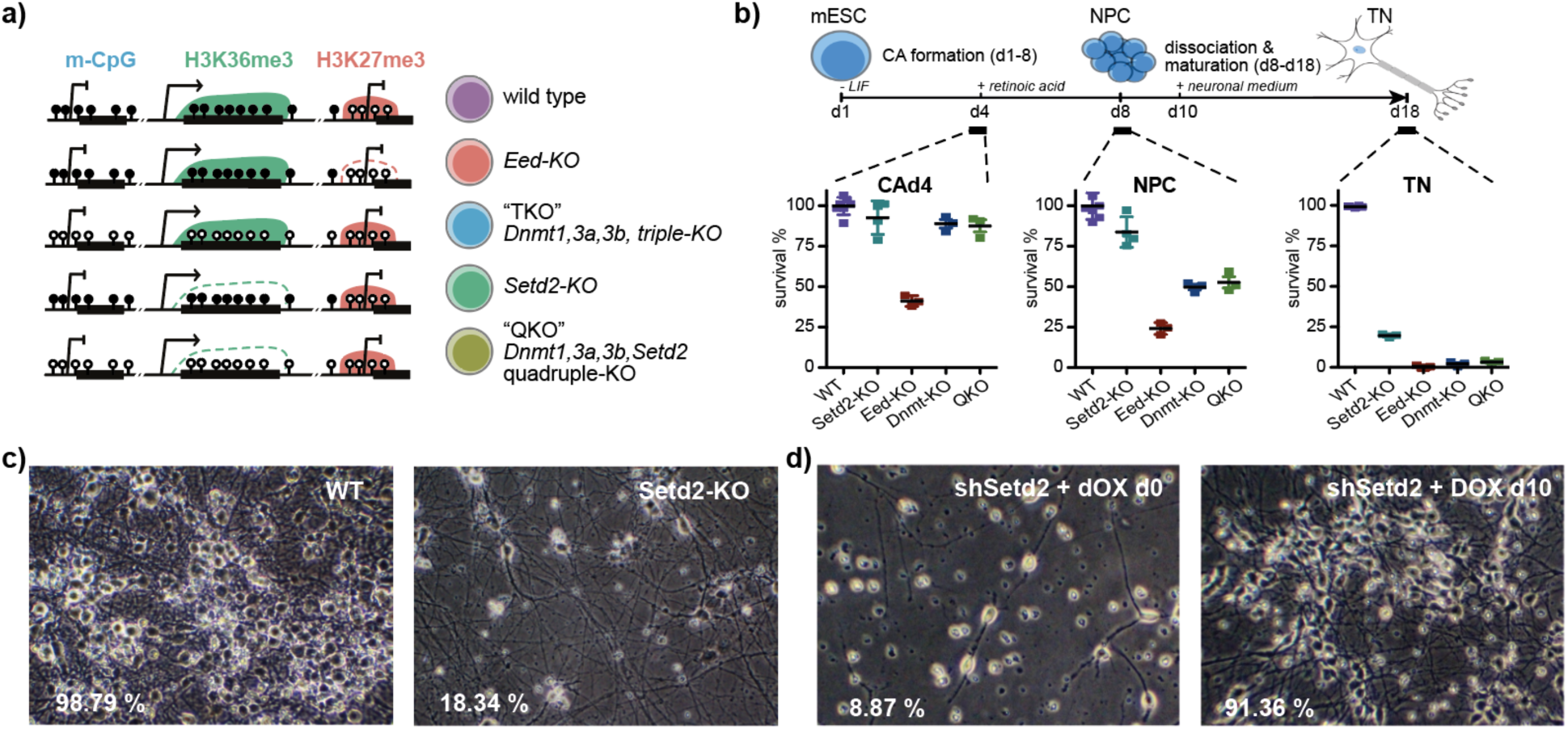
Stage-specific requirement for chromatin regulators during exit from pluripotency, lineage commitment, and terminal differentiation. **a)** Schematic overview of mESC lines utilized in this study. *Eed*-KO lack H3K27me3, *Dnmt*-TKO lack DNA methylation (5mC), *Setd2*-KO cells lack H3K36me3, and QKO (*Setd2*-KO in *Dnmt*-TKO) cells lack both H3K36me3 and 5mC. **b)** Top: schematics depicting the neuronal differentiation protocol. Bottom: cell count assay using live-dead stain at the cellular aggregate stage day 4 (CAd4), neural progenitor cells (NPC) and terminal neurons stage day 14 (TN). Depicted are percentages for survival in WT, *Setd2*-KO, *Eed*-KO, *Dmnt*-TKO and QKO cells, calibrated to 100% survival in WT cells. Data obtained from three to five independent replicates is shown. **c)** Microscopy images of WT and *Setd2*-KO TNs at day d14 at 100x magnification. Average percentages of survival after dissociation obtained from three independent experiments are indicated. **d)** Same as in (c) but using TNs from Tet-inducible *Setd2* knockdown cell lines that were treated with 1 μg/ml doxycycline (DOX) from d0 to d14, or from d10 to d14. Continuous knock-down of *Setd2* results in neuronal cell death, while knock-down in mature neurons does not influence survival.

In contrast, we observed normal progression to the progenitor stage in the *Setd2-*KO mESCs, with similar numbers of surviving cells and lack of morphological differences compared to wild-type cells (Figure 1b and Supplementary Figure 3a). However, upon further differentiation to mature neurons, we observed that the survival of *Setd2*-KO-derived neural progenitors was strongly impaired, with around 20% developing into viable post-mitotic neurons (Figure 1b-d). This was reproduced in two independent knock-out clones and one constitutive *Setd2* knock-down cell line (Supplementary Figure 3c-d).

### SETD2 is required for correct establishment of neuronal gene expression programs

Following these observations, we wanted to explore if the reduced generation of neurons in absence of SETD2 is due to impaired establishment or failure to maintain neuronal cell identity. We used a doxycycline-inducible sh*Setd2* mESC line to knock-down *Setd2* at different timepoints during the neuronal differentiation (Supplementary Figure 3e). Cells that were treated constantly with doxycycline (day d0) or from early neural progenitor stages on (day d6 or d8) showed a low number of surviving and differentiated neurons, similar to the phenotype observed in the knock-out cells (Figure 1d and Supplementary Figure 4a). In contrast, induction immediately after the early steps of neuronal differentiation (day d10) did not severely affect cell survival, resulting in 86-91 % surviving neurons (Figure 1d and Supplementary Figure 4a-b). Reduced levels of SETD2 in post-mitotic neurons after doxycycline-induced expression of shRNA at timepoint d10 was confirmed by RT-qPCR (Supplementary Figure 4c).

To obtain further insight into the contribution of SETD2 or H3K36me3 to the establishment of neuronal gene expression programs, we performed bulk RNA-seq in wild type and *Setd2-KO* neural progenitor cells (NPC) at day 8 of neuronal differentiation. Differential gene expression analysis at this early stage of neuronal commitment showed that 232 genes were upregulated, and 319 genes were downregulated in the *Setd2*-KO neural progenitor cells (Figure 2a and Supplementary Table 1). Gene set enrichment (GSEA) and gene ontology (GO) analysis showed high enrichment for neurodevelopmental-related processes in the down-regulated gene set, suggesting a deregulation of neuronal transcriptional programs already at this stage (Figure 2b and Supplementary Figure 5a-d). This was also confirmed by Motif Activity Response Analysis ^26^ which identified motifs preferentially bound by REST and SOX2 in the promoters of the downregulated genes (Figure 2c). Genes showing increased gene expression were mainly related to developmental processes, reproduction and transcription, translation, and metabolic processes, and coincided with E2F1 and MYC/MAX TF binding sites in their promoters, indicating increased proliferation and reduced differentiation (Figure 2b-c and Supplementary Figure 5a, b and d). Furthermore, we observe that long genes and genes with long transcripts were more likely to be downregulated in absence of SETD2 (Supplementary Figure 5e-f), which is also in accordance with the notion that neuronal-specific genes are longer than non-neuronal genes ^27,28^.

**Figure 2.**
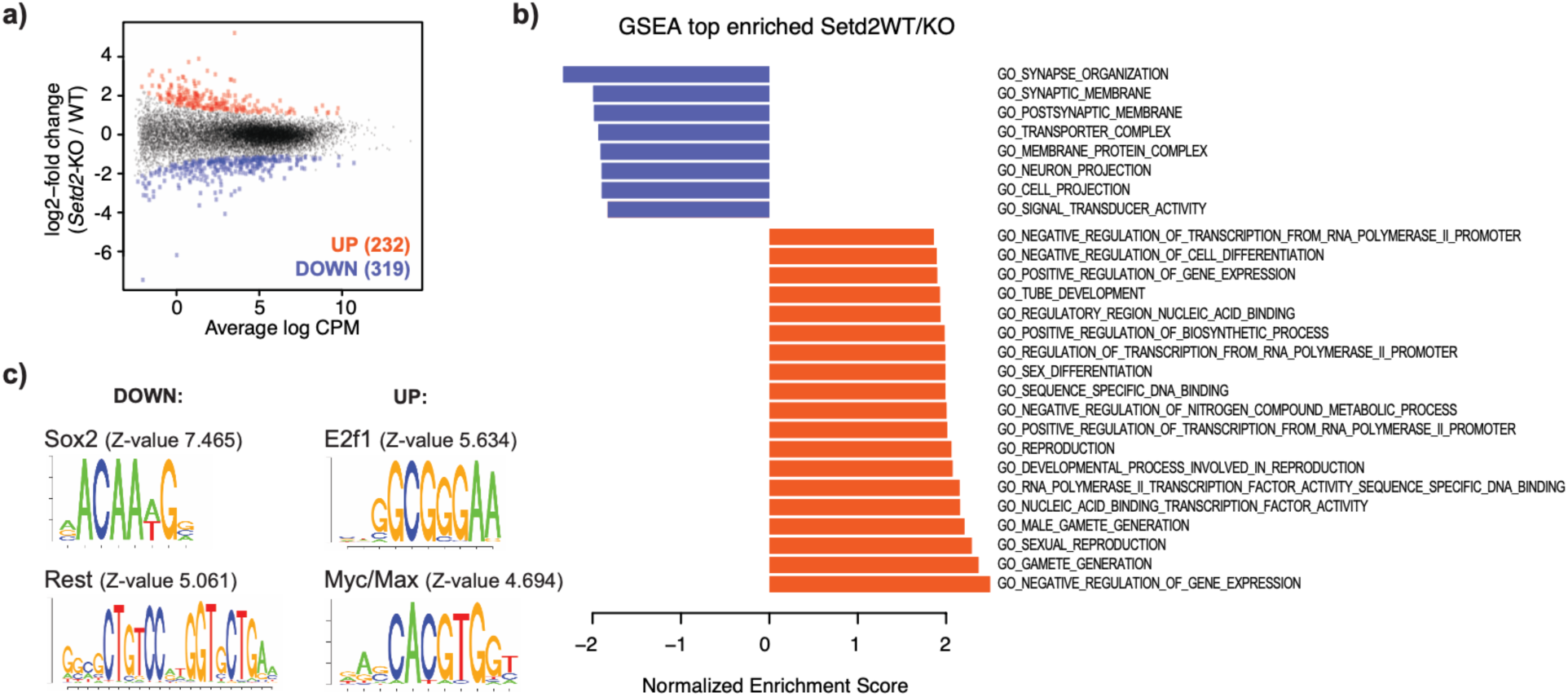
SETD2 is required for establishment of neuronal gene expression programs. **a)** MA plot of bulk PolyA-RNA-seq results depicting differences in gene expression between wild-type (WT) and *Setd2*-KO NPCs (p-value < 0.05, LFC > I1I). **b)** Gene set enrichment analysis (GSEA) for differentially expressed genes indicated a reduction of neuronal gene expression in *Setd2*-deficient NPCs. Shown are the top-most enriched terms. **c)** Motif Activity Response Analysis identifies transcription factor motifs preferentially bound by Rest and Sox2 in the promoters of genes downregulated in *Setd2*-KO NPCs, and motifs coinciding with E2f1 and Myc/Max TF binding in the promoters of upregulated genes.

### Catalytic activity of SETD2 and H3K36me3 are largely dispensable for neuronal differentiation

While the above results indicate that SETD2 is required for the successful initiation of neuronal gene expression programs, they do not explain how the observed phenotype is caused by absence of the SETD2 protein, loss of SETD2 catalytic activity, or loss of H3K36me3. To address this, we first performed genome-wide chromatin measurements in wild-type and *Setd2*-KO neural progenitors. Besides the global removal of H3K36me3 we did not observe drastic changes in chromatin modifications or chromatin accessibility in absence of SETD2 (Figure 3a and Supplementary Figure 6a-b). By focusing our attention on chromatin changes at promoters and gene bodies of genes with differential gene expression in absence of SETD2, we observed correlated changes in RNA Pol2 occupancy, histone acetylation, and chromatin accessibility, as expected (Supplementary Figure 6c-d). However, while we observe a modest correlation between increased histone acetylation and loss of H3K36me3 at gene bodies (Figure 3b) as previously reported in yeast ^8,29,30^, we do not observe any relationship between differentially expressed genes and loss of H3K36me3 at gene bodies (Figure 3c and Supplementary Figure 6d).

**Figure 3.**
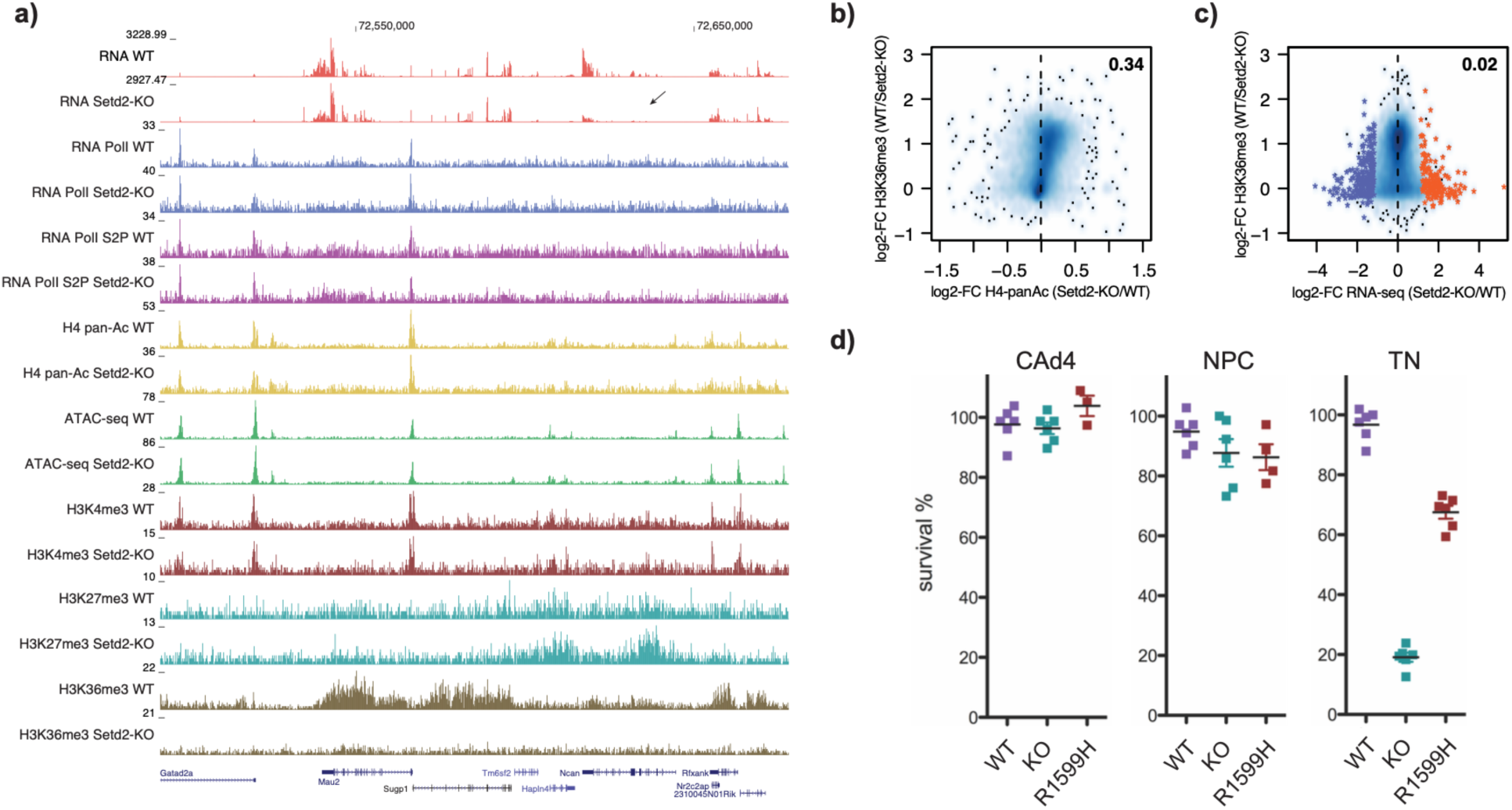
Catalytic activity of SETD2 is dispensable for neuronal differentiation. **a)** Representative genome browser view exemplifying differences in ChIP-seq signal of various chromatin marks and ATAC-seq signal between wild-type and *Setd2*-KO NPCs. Shown are read counts per 100 bp for ChIP-seq samples. Arrow exemplifies a downregulated neuronal gene *Ncan*. **b)** Scatterplot showing a minor correlation between the measured H4-panacetylation changes in *Setd2*-KO over WT NPCs and H3K36me3 levels at gene bodies measured in WT cells. **c)** Scatterplot showing changes in gene expression measured in *Setd2*-KO over WT NPCs and their relation to H3K36me3 levels at gene bodies measured in WT cells. **d)** Cell survival assay using live-dead stain at CAd4, NPC and TN stage (d14). Depicted are percentages for survival of WT, *Setd2*-KO and SET domain catalytic mutant (R1599H) cells.

This suggests an H3K36me3-independent role for SETD2 in establishing correct gene expression programs. To further test the requirement for the catalytic activity of SETD2 during neuronal differentiation, we engineered a point mutation in the catalytic SET domain (R1599H) ^31^ in both endogenous *Setd2* alleles (Supplemental Figure 6e-f). This point mutation resulted in global reduction of H3K36me3 (Supplemental Figure 6f). Surprisingly, despite the global absence of H3K36me3, we observe successful differentiation of neuronal cells in presence of this catalytically inactive SETD2 enzyme (∼ 65%, Figure 3d), supporting the notion that catalytic activity and H3K36me3 are largely dispensable for neuronal differentiation.

### Reduced association of the PAF1 complex with the elongating RNA Pol II in absence of SETD2

The results above suggest a catalytically-independent role of SETD2 required during the establishment of neuronal gene expression. To test how SETD2 influences transcription independent of H3K36me3, we investigated the protein interaction network of the elongating RNA Polymerase II in absence of SETD2. Towards this, we fused the SRI domain of SETD2, which is responsible for binding the serine-2-phosphorylated CTD of RNA Pol II ^32^, to the promiscuous biotin ligase TurboID ^33^. This enables the identification of proteins associated with the elongating RNA Pol II by proximity biotinylation. We stably expressed this fusion protein in wild type and *Setd2*-KO neural progenitor cells and performed biotin proximity-ligation followed by stringent protein enrichment as previously established ^23^ (Figure 4a and Supplemental Figure 7a). As a background control, we used an NLS-TuboID fusion protein lacking the SRI domain, expressed in the same cell lines (Supplemental Figure 7a). The enriched biotinylated proteins in wild type and *Setd2*-KO neural progenitors were detected by quantitative, label-free liquid chromatography tandem mass spectrometry (LC–MS/MS), resulting in an average of 890 proteins collectively detected in all samples (Supplemental Figure 7b). Since the background NLS-TurboID samples did not show significant differences between the two genetic backgrounds (Supplemental Figure 7c), we identified statistically-significant enriched proteins using a pooled background set. This resulted in 62 proteins significantly enriched in WT and 61 proteins enriched in *Setd2*-KO cells (Supplemental Figure 7d-f and Supplementary Table 2). In both samples, we identified proteins associated with GO Molecular Functions associated with RNA Pol II and RNA binding (Supplemental Figure 8a).

**Figure 4.**
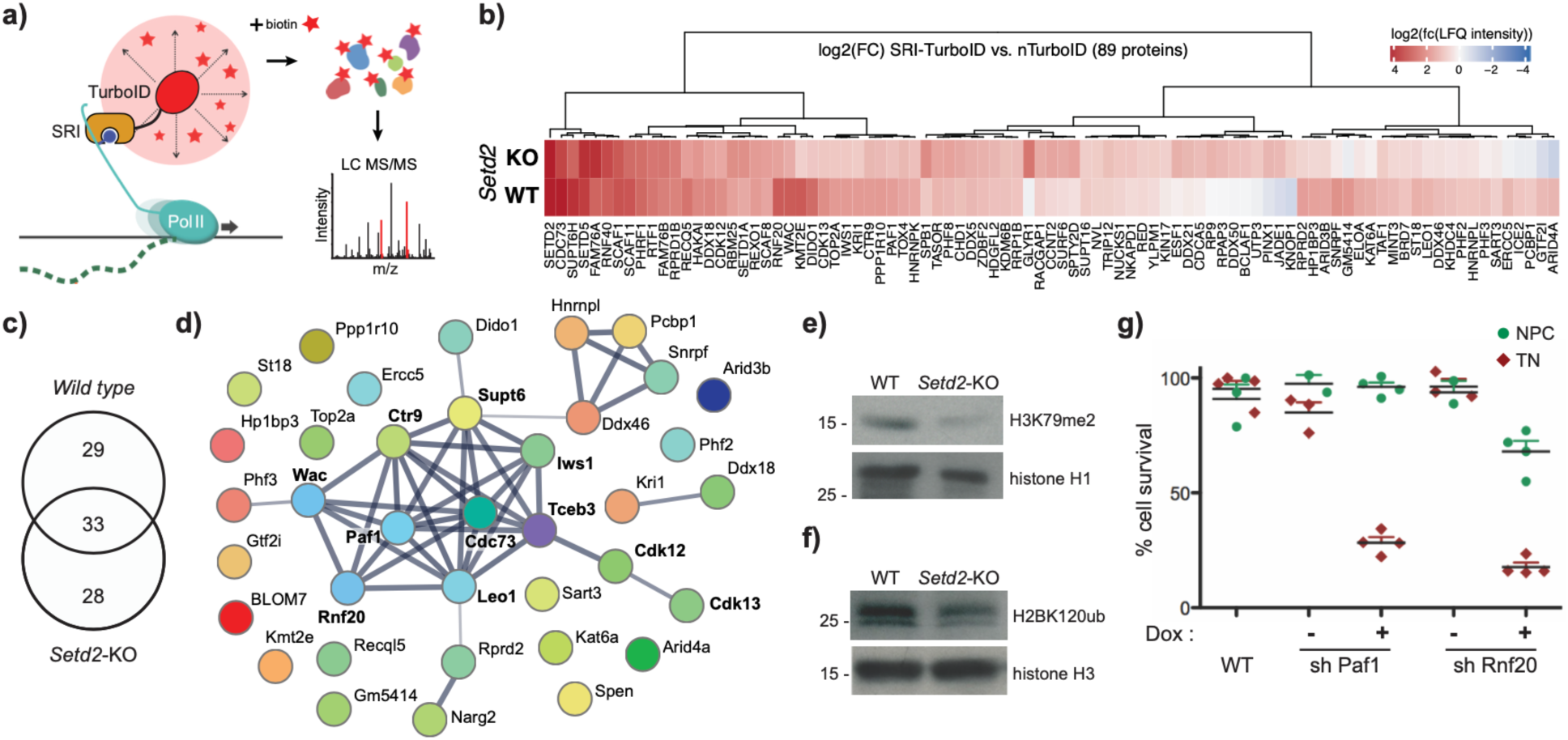
Loss of SETD2 results in reduced association of elongation factors with RNA Pol II. **a)** Schematic overview of the ChromID setup using SETD2-SRI domain fused to TurboID to detect protein-protein interactions on the elongating RNA Pol II in wild type and *Setd2*-KO NPCs using mass spectrometry. **b)** Heatmap indicating significantly enriched proteins in wild type and *Setd2*-KO NPCs over a background control expressing NLS-TurboID. Results obtained from four independent replicates. Colors indicate log2-fold change in LFQ intensity from SETD2-SRI over NLS-TurboID in the respective experiments. **c)** Venn diagram indicating the overlap between proteins significantly enriched by SRI-TurboID over nTurbo in Setd2-KO and wild type cells. (FDR-corrected two-tailed *t*-test: FDR = 0.01, s0 = 0.1, log_2_ FC > 0, *n* = 4 independent replicates). **d)** STRING analysis of proteins that show reduced RNA Pol II interactions in *Setd2*-KO NPCs identified factors involved in RNA Pol II elongation. **e-f)** Immunoblot analyses indicates reduced H3K79me2 and H2BK120Ub levels in *Setd2*-KO NPCs. Histones H1 and H3 served as loading controls. **g)** Cell survival assay using live-dead stain indicated reduced TN survival upon sh-mediated knock-down of Paf1 or Rnf20. Depicted are percentages for survival obtained from four independent replicates at NPC and TN stage.

Among the RNA Pol II-interacting proteins, we identified several factors that were depleted in absence of SETD2. These included proteins involved in transcription and histone acetylation, according to their GO Molecular Function (Figure 4b-d and Supplemental Figure 8b-c). Interestingly, several factors of the core PAF1 complex (PAF1, CDC73, LEO1, CTR9) and related proteins involved in histone H2B Lys-120 monoubiquitination (RNF20, WAC), as well as elongation-associated proteins (CDK12, SUPT6, TCEB3, IWS1, PHF3) were depleted in absence of SETD2 (Figure 4d). PAF1C plays a crucial role in transcriptional elongation and regulates the deposition of H2BK120ub (via RNF20/40) and H3K79me2 at transcribed genes ^34^. Indeed, we observe a reduction in global H3K79me2 levels, and minor reduction in H2BK120ub in absence of SETD2 (Figure 4e-f). These findings suggest that in absence of SETD2 the association of elongation factors with RNA Pol II could be diminished, resulting in transcriptional defects. To test if the identified changes in elongation factor interactions with the RNA Pol II would impact neuronal differentiation, we generated cell lines with dox-inducible shRNA expression against Paf1 and further Rnf20 (Supplementary Figure 8d). In both cases, we observe a strong effect on differentiation after reduction of RNF20 and PAF1 levels, with only 15-20% neuronal differentiation potential, and resembling the results obtained in absence of SETD2 (Figure 4g and Supplementary Figure 8e).

### Induced expression of neuronal master transcription factors can partially compensate absence of SETD2 during differentiation

The results above indicate that absence of SETD2 and the resulting destabilization of the PAF1 complex could influence the establishment of neuronal transcription, causing the observed differentiation failure. We argued that this deficit could be rescued by enforcing increased transcriptional initiation from neuronal genes. To test this, we generated *Setd2*-KO ES cell lines with *Tet*-inducible expression of the two bHLH transcription factors *Neurogenin 1* and *Neurogenin 2* (iNgn1/2) (Figure 5a and Supplementary Figure 9a). Induced expression of these factors has been shown to enable rapid neurogenesis by inducing genetic programs involved in the transition from stem cells to pre-natal neurons ^35^. We induced their expression in *Setd2*-KO at the NPC stage and evaluated their ability to rescue incorrect establishment of neuronal gene expression programs through measuring the success of neuronal differentiation. Induction of Ngn1/2 indeed increased the survival rate of *Setd2*-KO neurons from 15.87 % to 44.48 %, compared to 88.25 % in WT +dox (Figure 5b and Supplementary Figure 9b). In addition, RT-qPCR analysis for selected lineage marker genes confirmed this partial rescue (Figure 5c), suggesting that increasing the frequency of initiation from neuronal genes in presence of neuronal master regulators can partially overrule the necessity for SETD2.

**Figure 5.**
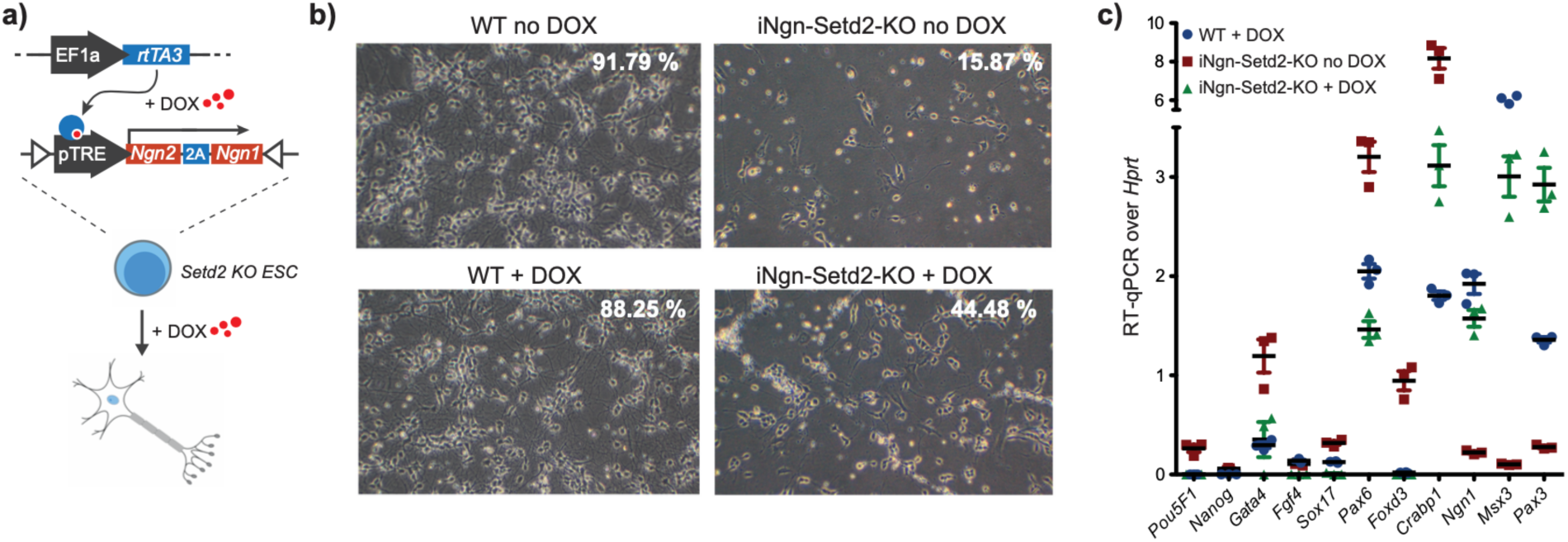
Overexpression of master transcription factors overrides SETD2 requirement for neuronal differentiation. **a)** Schematic overview of mESC line generation harboring a Tet-inducible Neurogenin2-Neurogenin1 (iNgn) master transcription factor cassette in a *Setd2*-KO background using recombination-mediated cassette exchange. **b)** Microscopy images of *in vitro-*derived neurons (d14) shows successful differentiation of neuronal cells derived from wild-type ESCs and partial rescue of cell death in *Setd2*-deficient cells by overexpression of an inducible Neurogenin2-Neurogenin1 fusion factor (i*Ngn*). Similar results were obtained from two independent replicates. Cells were treated with 1 μg/ml doxycycline (+DOX) during the entire *in vitro* differentiation, or with DMSO (no DOX). Mean neuronal survival rate (in percent) from three independent replicates is shown. **c)** RT-qPCR of various lineage marker in WT and *Setd2*-KO NPCs, the latter expressing Neurogenin1 and Neurogenin2 upon 1 µg/ml doxycycline treatment (n=3). Target genes: *Oct4* (*Pou5f1*), *Nanog* - embryonic stem cell markers; *Pax3*, *Pax6*, *Ngn1*, *Crabp1*, *Msx3*, *Foxd3* - neuronal marker; *Fgf4* - growth factor, *Sox17* - parietal endoderm marker. Gene expression was calibrated to *Hprt* - housekeeping gene expression.

## Discussion

By directly comparing the requirement for DNA methylation, H3K27me3 and H3K36me3 during exit from pluripotency and neuronal differentiation, we show their requirement for re-wiring of transcriptional programs at different stages of differentiation. Most-notably, and in contrast to DNA methylation and H3K27me3, SETD2 and H3K36me3 are dispensable for exit from pluripotency and neuronal lineage commitment. SETD2 however, is required during later stages of neuronal differentiation, where only 10-20% of the differentiated *Setd2*-KO neurons survive. This mirrors observations from KO experiments in mice, where *Setd2*-/- is embryonically lethal only at day E10.5, leading to growth defects including forebrain hypoplasia and unclosed neural tubes ^18^. By using an inducible system where we depleted SETD2 at different stages of neuronal differentiation, we further show that SETD2 is required during the terminal differentiation stage of NPCs, while removal of SETD2 in differentiated neurons did not influence their survival. This suggests that SETD2 could play a role during the establishment of neuronal gene expression programs rather than maintenance in differentiated neurons. This was further corroborated by transcriptional defects in neuronal gene expression that were already observed in *Setd2*-KO NPCs, despite showing no apparent phenotypes.

H3K36me3 has been proposed to play important roles in regulating transcriptional elongation, gene body chromatin, or to prevent spurious transcription initiation from gene bodies ^7^. By comparing genome-wide measurements for chromatin states and transcription in wild type and *Setd2*-KO NPCs, we were not able to associate the observed changes in gene expression with the loss of H3K36me3. At the same time, genes that lost H3K36me3 in absence of SETD2 did not display any changes in transcription or major differences in RNA Pol II occupancy. This lack of correlation points to a potential non-catalytic role of SETD2, which we further substantiated by showing that neurons expressing a catalytic dead SETD2 were able to differentiate. This finding is in line with a previous study, which showed that H3.3K36A mutations in both H3.3 genes are compatible with neuronal differentiation of ESCs and that the resulting changes in H3K36me3 do not necessary alter gene expression ^36^. Non-catalytic roles for histone modifying enzymes have been previously described, indicating that cellular functions of chromatin modifiers can extend beyond their histone-modifying capacity ^22^. For example, the catalytic activity of the H3K4me1 methyltransferases MLL3-MLL4 has been reported to be dispensable for naïve to primed transition of mESC, which is blocked in full KO cells ^37^.

To investigate the mechanism underlying the non-catalytic role of SETD2 during neuronal differentiation, we fused the SRI domain of SETD2 that binds to the Ser2-phosphorylated CTD of RNA Pol II ^32^ to the promiscuous biotin ligase TurboID in order to investigate the protein interactome of the elongating RNA Pol II in presence and absence of SETD2. Proximity biotinylation, followed by quantitative mass spectrometry, identified 90 significantly enriched proteins associated with RNA Pol II transcription and elongation machinery. Among these factors, we identified four out of five core subunits of the Polymerase Associated Factor 1 Complex (PAF1C; PAF1, CTR9, CDC73 and LEO1) and the PAF1C-associated factors SUPT6, IWS1, WAC, RNF20 to be depleted in *Setd2*-KO NPCs. PAF1C interacts with the elongating RNA Pol II in gene bodies and is essential for co-transcriptional processes and Pol II processivity ^38,39^. Absence of PAF1C results in decreased H3K36me3 ^40^ and H2B ubiquitination ^41^, further connecting the activity of this complex with histone modifications at transcribed genes. Along these lines, we also observe a reduction of RNF20, the E3 ligase responsible for H2B ubiquitination in mammals, as well as WAC, a factor involved in linking RNF20/40 with the elongating RNA Pol II ^42,43^, resulting in reduced H2BK120ub levels in *Setd2*-KO NPCs.

These results suggest that absence of SETD2 influences the association of PAF1C with the elongating polymerase, potentially attenuating productive transcriptional elongation. How SETD2 contributes to the association of PAF1C with the elongation machinery is unclear. One speculation could be that it directly contributes as a structural component that helps to stabilize PAF1C by interacting with the S2-phosphorylated Pol II CTD via it’s C-terminal SRI domain, while it’s N-terminal IDR, that has been identified to form condensates ^44^, could contribute to the stability of the elongation complex. Along these lines, PHF3, another S2-phosphorylated-CTD interacting protein, has been suggested to influence elongation through regulating phase separation of phosphorylated Pol II ^45^. Alternatively, SETD2 could influence PAF1C association with the elongating RNA Pol II indirectly. SUPT6 and IWS1 are additional factors that we have identified to be depleted from RNA Pol II in absence of SETD2. SUPT6 has been shown to facilitate recruitment of PAF1C to RNA Pol II and is required for PAF1C’s association with elongating RNA Pol II ^46,47^, while the SUPT6 partner protein, IWS1, has been suggested to connect SETD2 and SUPT6 ^48^. This could hint to an indirect role of SETD2 in PAF1C association with the active elongation complex through SUPT6. Future biochemical studies will allow to address the exact structural role of SETD2 in promoting PAF1C stability on the elongating machinery.

Importantly, conditional depletion of PAF1C members in mammalian cells results in decreased processivity, promoting poor elongation of RNA Pol II and reduction of full-length transcripts ^39,49^. This results in skewed RNA production from genes based on their length, where long genes are more affected compared to shorter genes, failing to complete transcription. Interestingly, neuronal genes tend to be very long ^27,28^ and we observe that in *Setd2*-KO NPCs, downregulated genes are generally longer indicating a potential link between reduced elongation efficiency of long transcripts and sub-optimal expression of genes required for neuronal differentiation. Finally, to see if we can overcome this elongation deficiency caused by SETD2 depletion, we forced increased expression of neuronal genes through inducing ectopic expression of the master transcription factors Neurogenin 1 and Neurogenin 2 in *Setd2*-KO NPCs. This led to the re-expression of several neuronal marker genes and increased generation of fully differentiated neurons indicating that the elongation deficiency can be compensated.

Taken together, we reveal a novel, non-catalytic role of SETD2 in promoting transcription through contributing to the association of the PAF1C complex with the elongating RNA Pol II. Although the PAF1C interaction with RNA Pol II is only mildly affected in absence of SETD2, this is likely sufficient to influence the productive transcription of long neuronal genes, therefore impacting neuronal differentiation potential. *Setd2* loss-of-function mutations or deletions are frequent in renal cell carcinoma and associated with increased metastasis ^50,51^. In addition, heterozygous mutations in *Setd2* cause developmental delay, intellectual disability, brain deformities, and macrocephaly ^52–54^. Our study identifies the involvement of SETD2 in promoting neuronal gene expression through stabilizing the RNA Pol II elongation machinery, offering novel insights into potential mechanisms underlying these diseases.

## Materials and methods

### Cell line generation, cell culture, and neuronal differentiation

Mouse embryonic stem cells (HA36CB1) were cultured on 0.2 % gelatine-coated dishes in DMEM (Invitrogen), supplemented with 15 % fetal calf serum (Invitrogen), 1x non-essential amino acids (Invitrogen), 1x Glutamax (Invitrogen), homemade leukemia inhibitory factor (LIF), and 0.001 % b-mercaptoethanol (Invitrogen) at 37°C and 7 % CO2. ESC lines were differentiated as previously described ^25^, except that no feeder cells were used. Microscopy images were taken at 100 x magnification. Cell count assays were performed using live/dead stain and TC20™ Automated Cell Counter (BioRad).

*Setd2*-KO in *Dnmt*-triple KO mouse ESC background was generated using CRISPR/Cas9. The Cas9 sgRNA sequences as previously described ^12^, were cloned into the pX330-U6-Chimeric_BB-CBh-hSpCas9 (Addgene 42230). Transfections together with pRR-Puro recombination reporter (Addgene 65853) were conducted using Lipofectamine 3000 reagent (L3000015, Thermo Fisher Scientific) at a 2:1 Lipofectamine/DNA ratio in OptiMEM (31985070, Thermo Fisher Scientific). 36 hours later cells were selected with 2 μg/ml puromycin for another 36 hours. ESCs containing a homozygous CGT to CAC in-frame mutation at position R1599H of *Setd2* were generated using CRISPR/Cas9. Here, donor plasmids for homologous repair, containing 1kb homologous sequences upstream and downstream of the cut site and a mutated PAM site, were co-transfected in a 1:2.5 (sgRNA:donor) ratio together with sgRNA and the pRR-Puro reporter using lipofection. The R1599H mutation was confirmed by PCR and sanger sequencing, as well as immunoblotting for H3K36me3 levels.

*Setd2*-KO ESCs with inducible expression of Neurogenin 1 and 2 (iNgn1/2) were generated as described previously with adaptations ^35^. A doxycycline-inducible rtTA3 system (Addgene 61472) was randomly integrated using 20 µg of linearised plasmid with bleomycin resistance. Cells were treated with 200 µg/ml Zeocin (InvivoGen). The TetON-inducible Ngn2-2A-Ngn1 ESCs in *Setd2*-KO background were then obtained by recombinase-mediated cassette exchange (RMCE) ^55^ with an expression plasmid for CRE recombinase in a 1:0.6 DNA ratio. Similarly, shRNA-inducible ESCs were generated using a TetON-inducible system for a shRNA against *Setd2*, expressed from the RMCE site. Positive clones were confirmed by Sanger sequencing, RT-qPCR and immunoblotting. Oligo sequences available in Supplementary Table 3. Inducible shRNA-mediated knockdown as well as iNgn1/2 cell lines were treated with 1 µg/ml doxycycline (Sigma Aldrich).

### Surface marker and cell cycle analysis using flow cytometry

Single-cell suspensions were obtained through trypsinization and filtered through 40-um cell strainers (BD Biosciences). For cell cycle and Ki-67 measurements, single-cell suspensions were fixed and permeabilised for 30 min at 4°C with Foxp3/Transcription Factor Staining Buffer set (eBioscience). Anti-Ki-67 or isotype control (eBioscience) was added and incubated for 45 min at RT in permeabilisation buffer. DAPI (5 μg/ml, Sigma) was added as a fluorescent DNA stain 5 min prior to FACS measurements and incubated at RT in the dark. For live dead cell exclusion, LIVE/DEAD Fixable Near-IR Dead Cell Stain (Invitrogen) was used. Samples were acquired and data was analysed as described in ref. ^23^.

### Protein extraction and immunoblotting

Crude nuclear extracts cells were obtained as described in ref. ^56^, histones were acid extracted according to ref. ^23^. Membranes were blocked with 5 % milk in TBS with 0.1 % Tween20 and incubated with primary antibodies against SETD2 (1:1000, A3194, ABclonal, LOT 0071240201), H3K36me3 (1:5000, ab9050, abcam, LOT GR3210075-1), H3K79me2 (1:2000, 04-835, milipore, clone NL-59), H2BK120ub (1:2000, MM-0029-P, Medimabs, clone NRO3), anti-histone H1 (1:5000, AE04, Millipore, LOT 3087175), anti-NEUROG1 (1:1000, sc-100332, Santa Cruz Biotechnology, LOT F1119), anti-LAMIN B1 (1:1000, sc-374015, Santa Cruz Biotechnology, LOT J3019), anti-HA (1:1000, ab9110, abcam), Streptavidin-HRP (1:20000, pierce, clone 21130), overnight at 4°C. Protein detection was facilitated using species-specific antibodies conjugated to horseradish peroxidase and Pierce® Peroxidase IHC Detection Kit (Thermo Scientific).

### RNA isolation, cDNA synthesis and RT-qPCR

RNA was isolated from mouse ESCs, cellular aggregates, NPCs und terminal neurons using the RNeasy Plus mini kit (Qiagen). Coding DNA was synthesized from 2 μg isolated RNA with SuperScript III First-Strand Synthesis (Invitrogen) for 60 min at 50°C using Oligo(dT) (Thermo Fisher), followed by heat-inactivation for 10 min at 85°C. Residual RNA was digested with 2 units RNaseH (NEB) for 20 min at 37°C. Target sequences were quantified by real-time qPCR analysis using a KAPA SYBR FAST qPCR Kit (Sigma Aldrich) on a QuantStudio 5 Real-Time PCR System (Applied Biosystems). Comparative quantification (ddCt) was used to determine transcript levels relative to the housekeeping gene *Hprt*. Each sample was run in at least technical triplicates. Oligo sequences are available in Supplementary Table 3.

### PolyA RNA-sequencing and differential gene expression analysis

Total RNA was isolated from NPCs using the RNeasy Plus mini kit (Qiagen). RNA integrity was measured using a model 2100 Bioanalyzer (Agilent). PolyA-tailed mRNAs were isolated and enriched using NEB Next Poly(A) mRNA Magnetic Isolation Module according to manufacturer’s instructions. Libraries for 1 µg mRNA were prepared using NEB Next UltraTM II Directional RNA Library Prep Kit for Illumina. Sequencing of library pools and read processing were performed on Illumina NovaSeq according to Illumina standards, with 150-bp single-end sequencing. Sequencing reads were trimmed using trimgalore to remove adapter sequenced and aligned using STAR ^57^ using standard options based on the gene transcript annotation gencode.mouse.v1.annotation.gtf (NCBIM37, mm9). Gene counts were obtained using qCount() in QuasR ^58^ and differential gene expression was performed using the edgeR package with significance set to p-value < 0.05 and log fold change > I1I ^59^. Gene ontology enrichment analysis on differentially expressed was performed using the goana() function in edgeR. GSEA analysis was performed using FGSEA ^60^, while collapsing similar pathways for better representation. Motif response analysis was performed using the ISMARA online tool ^26^.

### Chromatin immunoprecipitation (ChIP) and ATAC -sequencing and read processing

Histone ChIP experiments and sequencing were performed as previously described {Villasenor:2020eh}. Here, 100 µg chromatin were incubated with 5 µg of either H3K36me3 (ab9050, abcam), total Pol II (MABI0601, MBL), Ser2-P RNA Pol II (ab5095, abcam), H3K4me3 (ab8580, abcam), H4-panacetyl (B_2687872, ActiveMotif), or H3K27me3 (C15410195, diagenode) antibody. Sequencing of library pools was performed on Illumina NovaSeq according to Illumina standards, with 150-bp single-end sequencing. The ATAC-seq reaction was performed with 50,000 cells as previously described in ref. ^61^ with minor adjustments of the protocol, using the Nextera DNA Library Prep Kit (Illumina) together with the barcoded primers from the Nextera Index Kit (Illumina). In brief, an additional size selection step was performed after the first 5 cycles of library amplification. For this, the PCR reaction was incubated with 0.6 x volume of Ampure XP beads (Beckman Coulter) for 5 min to allow binding of high molecular weight fragments. Beads containing long DNA fragments were separated on a magnet and the supernatant containing only small DNA fragments below roughly 800 bp were cleaned up using MinElute PCR purification columns (Qiagen). All libraries were amplified for 12 cycles in total, visualised and quantified with a TapeStation2200 (Agilent), and sequenced as mentioned above.

Library demultiplexing was performed following Illumina standards. Samples were filtered for low-quality reads as well as adaptor sequences and mapped to the mouse genome (NCBI Build 37 mm9, July 2007) using the BOWTIE1 algorithm allowing for two mismatches and only unique mappers were used. Identical reads from PCR duplicates were filtered out. Wiggle tracks were obtained with QuasR and visualised with the UCSC genome browser. For genomic analyses, read counts overlapping with promoters (2 kb around TSS) and gene bodies (+2 kb downstream from TSS) were used.

### ChromID and label-free MS data acquisition and analysis

ChromID samples were prepared as previously described in ref. ^23^. In brief, nuclear extraction of SRI-TurboID NPCs of wild-type and Setd2-KO background was performed after biotin incubation for 12h, followed by affinity purification through streptavidin beads, high-stringency washes, and on-bead digestion. MS data acquisition was performed as described in ref. ^23^. In brief, samples (quadruplicates per condition) were cleaned up by C18 StageTips. Peptides were detected by data-dependent acquisition via mass spectrometry. Proteins were identified and quantified from the raw acquisition data as well as processed using MaxQuant ^62^. The mouse reference proteome (UniProtKB/Swiss-Prot) version 2018_12 combined with manually annotated contaminant proteins was searched with protein and peptide false discovery rate (FDR) values set to 1% and the match-between-runs algorithm was enabled. Statistical analysis was conducted using Proteus^63^. LFQ intensity values were log2-transformed, and outlier samples determined based on low peptide or protein counts. Subsequently, Proteus’ limma-wrapper ^64^ was used to determine potential interactors in respective contrasts (bait vs. control of same genetic background) with an FDR of 0.05 as significance threshold. For gene ontology term analysis, differentially enriched proteins of bait conditions were combined and parsed to Enrichr ^65^.

## Acknowledgements

We thank members of the Baubec laboratory for their input and criticism. Furthermore, we thank members of the Functional Genomics Centre Zurich for their genomics and proteomics support. This work was supported by the Swiss National Science Foundation through SNSF Professorship (183722) and SNSF Sinergia (180354), by the European Research Council (865094 -ChromatinLEGO - ERC-2019-COG), and the EMBO Young Investigator program. C.A. acknowledges support from the UZH Candoc Grant and N.S. acknowledges support from EMBO postdoctoral fellowship and SNSF Ambizione grant (186012).

## Author contributions

C.A., R.P., S.B., X.B., and N.S. performed all experiments and generated reagents. C.A., R.P., D.R., and T.B. analyzed data and developed tools for data analysis. C.A. and T.B. wrote the manuscript with input from all authors.

