## Supplemental Figures and Information for "SETD2 promotes PAF1C interactions with the elongating RNA Pol II and is required for neuronal differentiation"

**Contents**

Supplementary Figures and Legends 1-9

Supplementary Table Legends 1-3

### Supplementary Figures

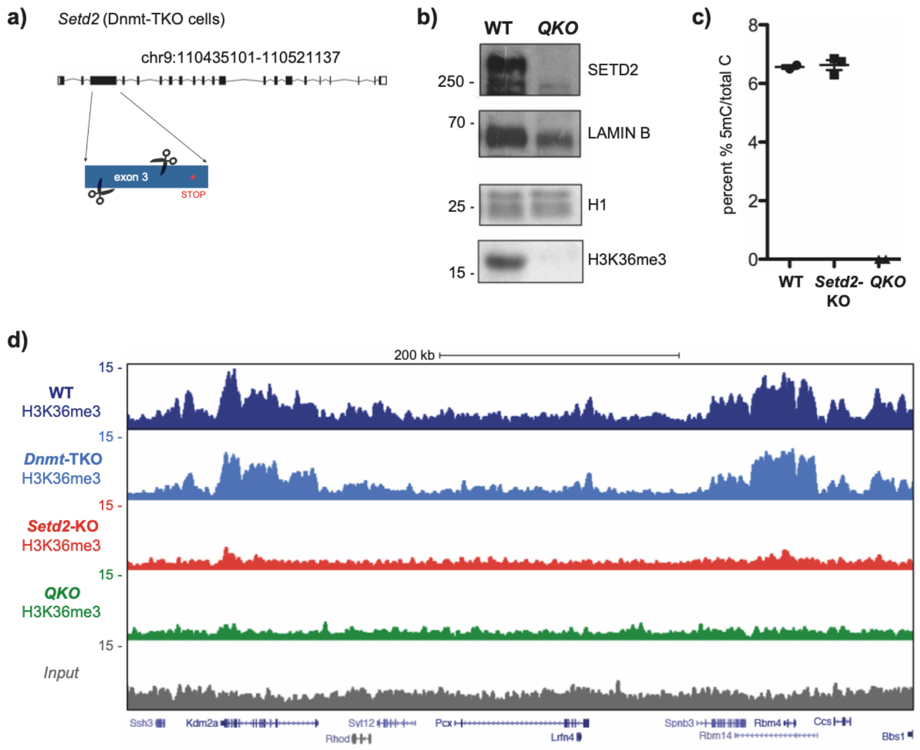


**Figure S1. *Setd2*-KO in *Dnmt1,3a,3b*-triple-KO mESC leads to loss of H3K36me3.**

**a)** CRISPR-Cas9 knock-out strategy for *Setd2* in *Dmnt*-TKO background (QKO). Two guide RNAs target exon 3, resulting in an out-of-frame deletion and a downstream premature stop codon. **b)** Immunoblot analysis for SETD2 and H3K36me3 levels in nuclear (top) and histone (bottom) extracts of wild-type (WT) and QKO mESCs. LAMIN B and H1 serve as loading controls. **c)** HPLC-MS measurement of methylcytosine to cytosine indicates absence of methylation in the QKO. Error bars denote standard deviation from three independent replicate measurements. **d)** Representative genome browser view of a chromosome 19 locus exemplifying differences in H3K36me3 signals between wild-type, *Dnmt* TKO*, Setd2*-KO, and QKO mESCs. Shown are read counts per 100 bp for ChIP-seq and input samples.

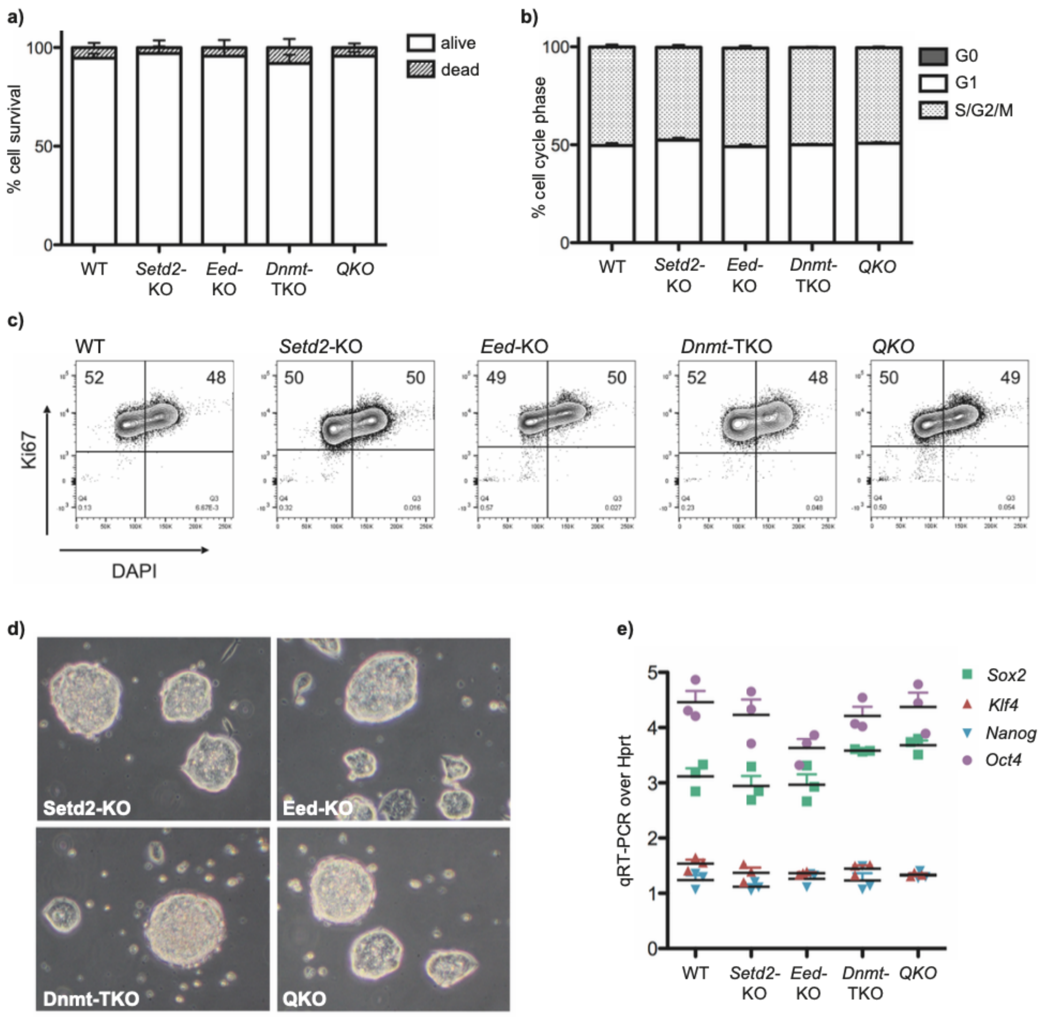


**Figure S2. Loss of chromatin marks does not impair survival, self-renewal capacity, proliferation, and transcription in mESCs.**

**a)** Wild-type (WT), *Setd2*-KO, *Eed*-KO, *Dnmt*-TKO and QKO mESCs were cultured in media containing leukemia-inhibitory factor (LIF) over three passages and dead/live cells were counted. **b)** Ki-67 and DAPI cell cycle analysis by FACS. Shown are percentages of the different phases of WT, *Setd2*-KO, *Eed*-KO, *Dnmt*-TKO and QKO ESCs, based on two independent replicates. **c)** Exemplary FACS analysis for Ki-67 and DAPI levels in WT, *Setd2*-KO, *Eed*-KO, *Dnmt*-TKO and QKO ESCs cells. Highlighted are percentages of cells in Ki67+ (G1 phase) and Ki67+/DAPI+ (S/G2/M phase) clusters. **d)** Microscopy images of *Setd2*-KO, *Eed*-KO, *Dnmt*-TKO and QKO mESCs in feeder-free culture at 100x magnification. **e)** RT qPCR detection of pluripotency marker genes *Nanog*, *Pou5F1* (*Oct4*), *Klf4*, and *Sox2 in* WT, *Setd2*-KO, *Eed*-KO, *Dnmt*-TKO and QKO mESCs. *Hprt* served as internal control.

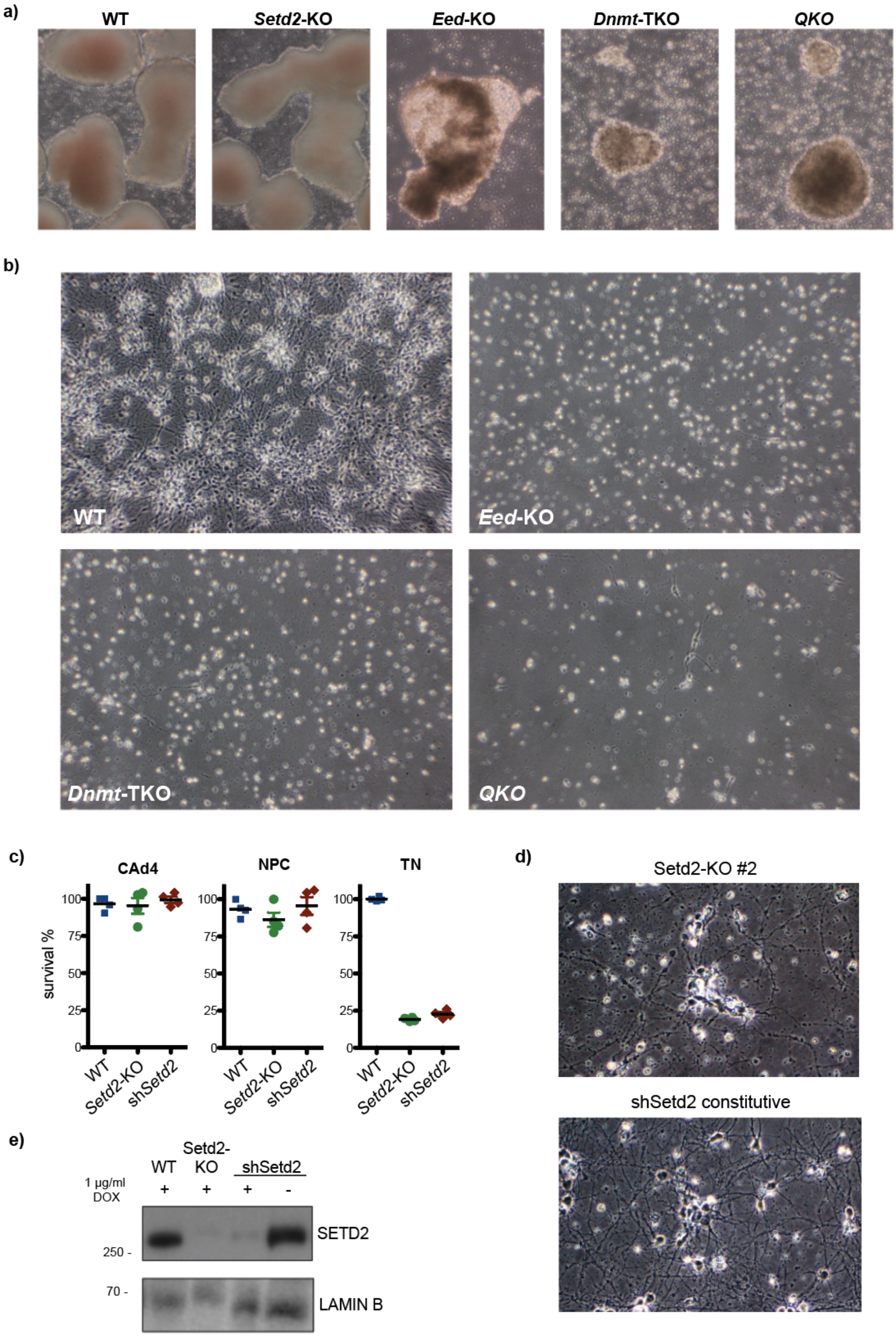
a)
WT
Setd2-KO
Eed-KO
Dnmt-TKO
QKO
b)
WT
Eed-KO
Dnmt-TKO
QKO
c)
CAd4
NPC
TN
survival %
100
75
50
25
0
WT
Setd2-KO
shSetd2
d)
Setd2-KO #2
shSetd2 constitutive
e)
WT
Setd2-
KO
shSetd2
1 µg/ml
DOX
+
+
+
-
SETD2
250 -
70 -
LAMIN B

**Figure S3. *Setd2*-KO, *Eed*-KO, *Dmnt*-TKO and QKO display loss of differentiation potential at different stages.**

**a)** Exemplary microscopy images of in vitro derived cellular aggregates at day 4 (CAd4) for WT, *Setd2*-KO, *Eed*-KO, *Dnmt*-TKO and QKO cells at 100x magnification. **b)** Exemplary microscopy images of in vitro derived, terminal neurons of wild-type, *Eed*-KO, *Dnmt*-TKO and QKO cells at day 14 at 100x magnification. **c)** Cell count assay using live-dead stain at cellular aggregate stage day 4 (CAd4), neural progenitor cells (NPCs), and terminal neurons (TNs) stage day 14. Depicted are percentages for survival in WT, *Setd2*-KO, and cells harboring a constitutively expressed shRNA against *Setd2* (sh*Setd2*). **d)** Exemplary microscopy images of *in vitro*-derived, terminal neurons of a second independently derived *Setd2*-KO clone and cells harboring a constitutively expressed shRNA against *Setd2* at 100x magnification. **e)** Immunoblot analysis for SETD2 levels in nuclear extracts of WT, *Setd2*-KO and Tet-inducible shSetd2 knockdown in terminal neurons at day d14 (+/- 1 µg/ml DOX). LAMIN B serves as loading control.

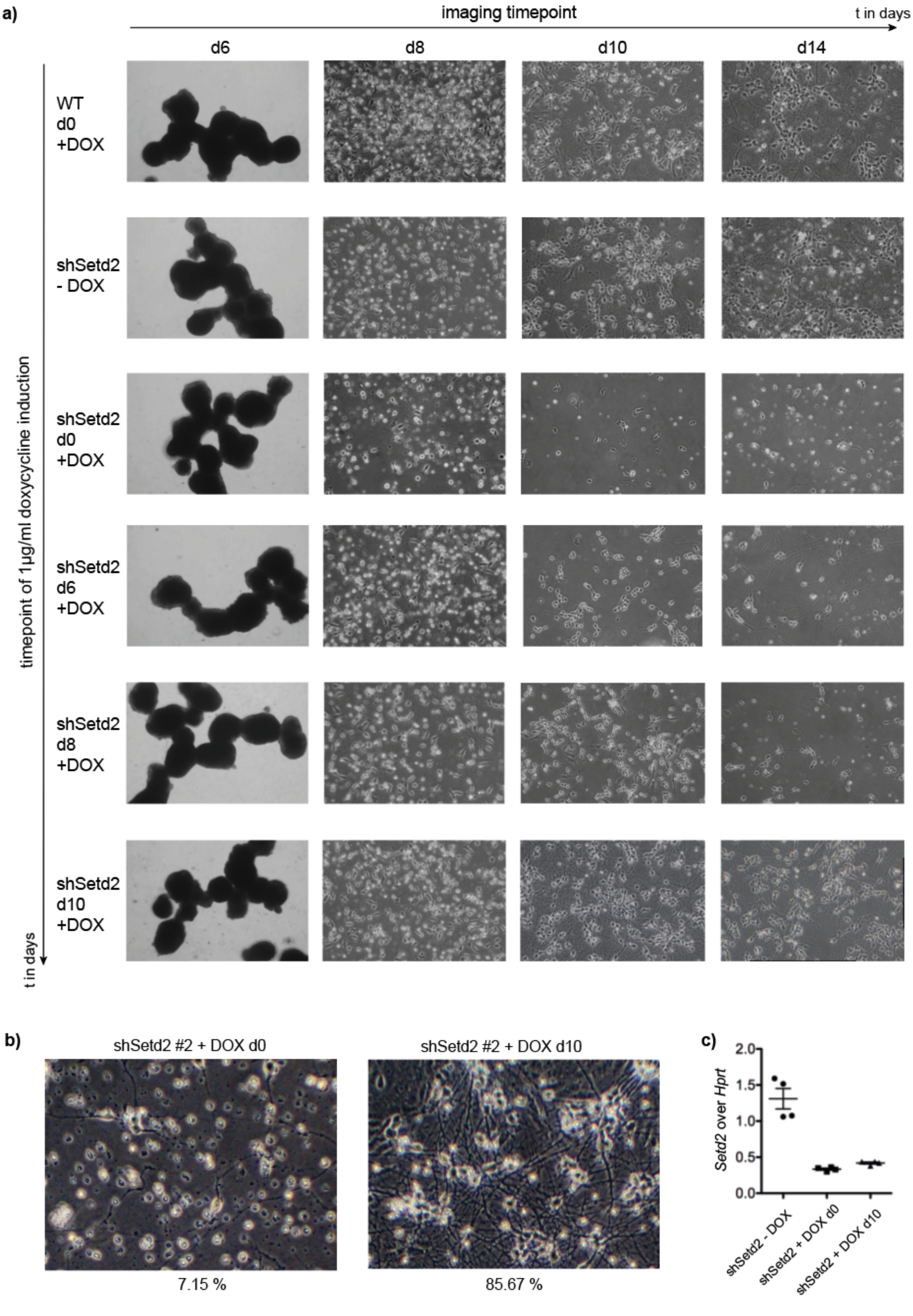
a)
imaging timepoint
t in days
d6
d8
d10
d14
timepoint of 1µg/ml doxycycline induction
t in days
WT d0 +DOX
shSetd2 - DOX
shSetd2 d0 +DOX
shSetd2 d6 +DOX
shSetd2 d8 +DOX
shSetd2 d10 +DOX
b)
shSetd2 #2 + DOX d0
shSetd2 #2 + DOX d10
7.15 %
85.67 %
c)
Setd2 over Hprt
2.0
1.5
1.0
0.5
0.0
shSetd2 - DOX
shSetd2 + DOX d0
shSetd2 + DOX d10

**Figure S4. Timed knock-down of *Setd2* reveals a requirement during early neuronal differentiation.**

**a)** Time-resolved microscopy images of *in vitro* differentiation using Tet-inducible *Setd2* knockdown cell lines. Shown are images starting from cellular aggregates (CA) day 6 over neural progenitors at d8 to terminal neurons at day 10 and 14. Treatment with 1 µg/ml doxycycline (DOX) was initiated at day 0, 6, 8, and 9; 100x magnification. **b)** Microscopy images of vitro-derived neurons at day d14 depicting a second, independently derived clone expressing a Tet-inducible shRNA against *Setd2*. Shown are average percentages of survival after dissociation (plated/attached) of three independent experiments; 100x magnification. Cells were treated with 1 µg/ml doxycycline (DOX) from the start of the differentiation (d0) or from day d10 on. **c)** RT qPCR of *Setd2* in Tet-inducible *Setd2* knockdown cells, differentiated to terminal neurons (day 14) shows efficient knock-down. Cells were treated with 1 µg/ml doxycycline, always or from day 10 on. *Hprt* served as internal control.

**a)**

*Setd2*-WT top GO signature

| | Term | Ont | N | DE | P.DE |
|---|---|---|---|---|---|
| GO:0007399 | nervous system development | BP | 2257 | 95 | 5.408119e-26 |
| GO:0097458 | neuron part | CC | 1920 | 85 | 1.659946e-24 |
| GO:0044456 | synapse part | CC | 1070 | 60 | 4.377557e-22 |
| GO:0045202 | synapse | CC | 1386 | 68 | 7.945682e-22 |
| GO:0048856 | anatomical structure development | BP | 5712 | 151 | 1.628307e-21 |
| GO:0007275 | multicellular organism development | BP | 5227 | 143 | 1.786169e-21 |
| GO:0048731 | system development | BP | 4638 | 133 | 1.903633e-21 |
| GO:0022008 | neurogenesis | BP | 1712 | 75 | 3.305885e-21 |
| GO:0048699 | generation of neurons | BP | 1613 | 71 | 3.742730e-20 |
| GO:0032502 | developmental process | BP | 6097 | 154 | 4.939505e-20 |
| GO:0030182 | neuron differentiation | BP | 1447 | 66 | 1.667239e-19 |
| GO:0098794 | postsynapse | CC | 718 | 46 | 3.001488e-19 |
| GO:0050808 | synapse organization | BP | 424 | 35 | 3.161904e-18 |
| GO:0032501 | multicellular organismal process | BP | 8064 | 177 | 3.015872e-17 |
| GO:0043005 | neuron projection | CC | 1478 | 63 | 3.594572e-17 |
| GO:0071944 | cell periphery | CC | 5266 | 134 | 5.432891e-17 |
| GO:0044459 | plasma membrane part | CC | 2926 | 93 | 5.524273e-17 |
| GO:0005886 | plasma membrane | CC | 5130 | 131 | 1.198679e-16 |
| GO:0030154 | cell differentiation | BP | 4125 | 114 | 1.333546e-16 |
| GO:0031175 | neuron projection development | BP | 1046 | 51 | 2.829015e-16 |

**b)**

*Setd2*-KO top GO signature

| | Term | Ont | N | DE | P.DE |
|---|---|---|---|---|---|
| GO:0048856 | anatomical structure development | BP | 5712 | 118 | 7.885580e-21 |
| GO:0007275 | multicellular organism development | BP | 5227 | 109 | 4.615013e-19 |
| GO:0032502 | developmental process | BP | 6097 | 118 | 1.991503e-18 |
| GO:0048513 | animal organ development | BP | 3430 | 84 | 4.710159e-18 |
| GO:0048731 | system development | BP | 4638 | 99 | 1.379404e-17 |
| GO:0009888 | tissue development | BP | 1899 | 60 | 1.611083e-17 |
| GO:0009653 | anatomical structure morphogenesis | BP | 2652 | 68 | 2.774195e-15 |
| GO:0009887 | animal organ morphogenesis | BP | 1048 | 41 | 5.012804e-15 |
| GO:0060429 | epithelium development | BP | 1160 | 42 | 3.109982e-14 |
| GO:0003700 | DNA-binding transcription factor activity | MF | 1023 | 38 | 2.851829e-13 |
| GO:0000981 | DNA-binding transcription factor activity, RNA polymerase II-specific | MF | 795 | 33 | 6.320282e-13 |
| GO:0032501 | multicellular organismal process | BP | 8064 | 125 | 6.466511e-12 |
| GO:0005488 | binding | MF | 13121 | 171 | 1.478950e-11 |
| GO:0043565 | sequence-specific DNA binding | MF | 1099 | 36 | 4.680072e-11 |
| GO:0030154 | cell differentiation | BP | 4125 | 79 | 4.696846e-11 |
| GO:0140110 | transcription regulator activity | MF | 1336 | 40 | 5.187143e-11 |
| GO:0035295 | tube development | BP | 1105 | 36 | 5.445985e-11 |
| GO:0042221 | response to chemical | BP | 4043 | 77 | 1.243958e-10 |
| GO:0048869 | cellular developmental process | BP | 4334 | 80 | 2.099922e-10 |
| GO:0009790 | embryo development | BP | 1186 | 36 | 3.799764e-10 |

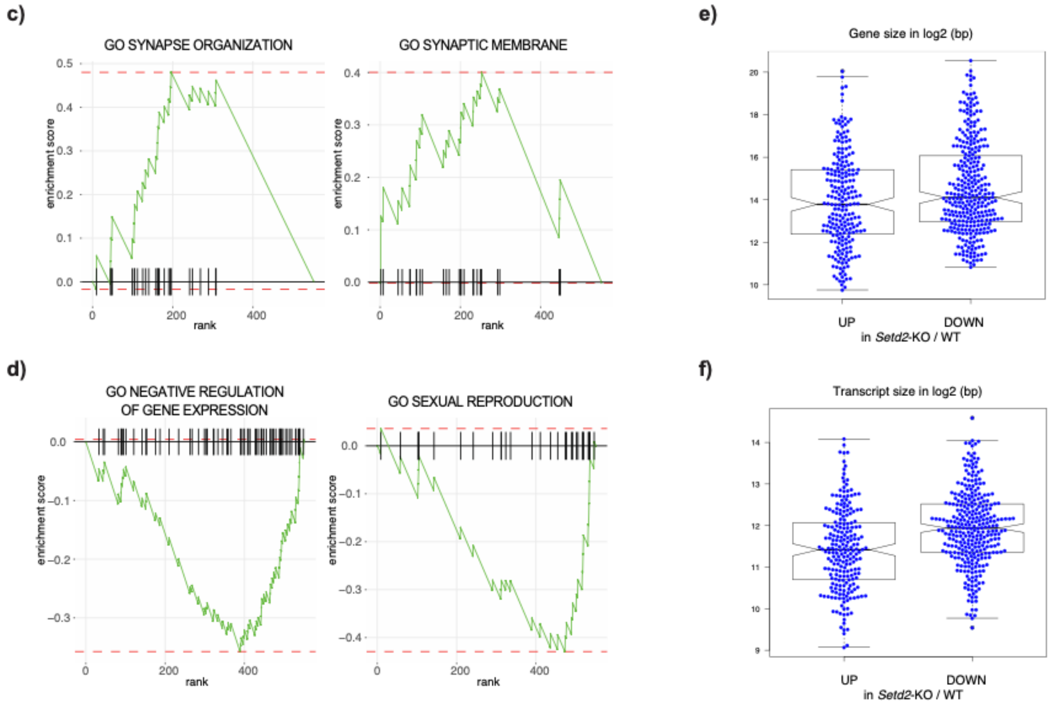


**Figure S5. Neuronal gene expression programs are disturbed in absence of SETD2/H3K36me3.**

**a-b)** Gene ontology (GO) term analysis for differentially expressed genes in WT and *Setd2*-deficient NPCs. Shown are the top-most enriched terms. **c-d)** Top two GO term enrichment scores for (C) downregulated and (D) upregulated genes in *Setd2*-KO cells. **e-f)** Boxplots showing gene size (E) and transcript length (F) for up- and downregulated genes in *Setd2*-KO versus wild-type NPCs.

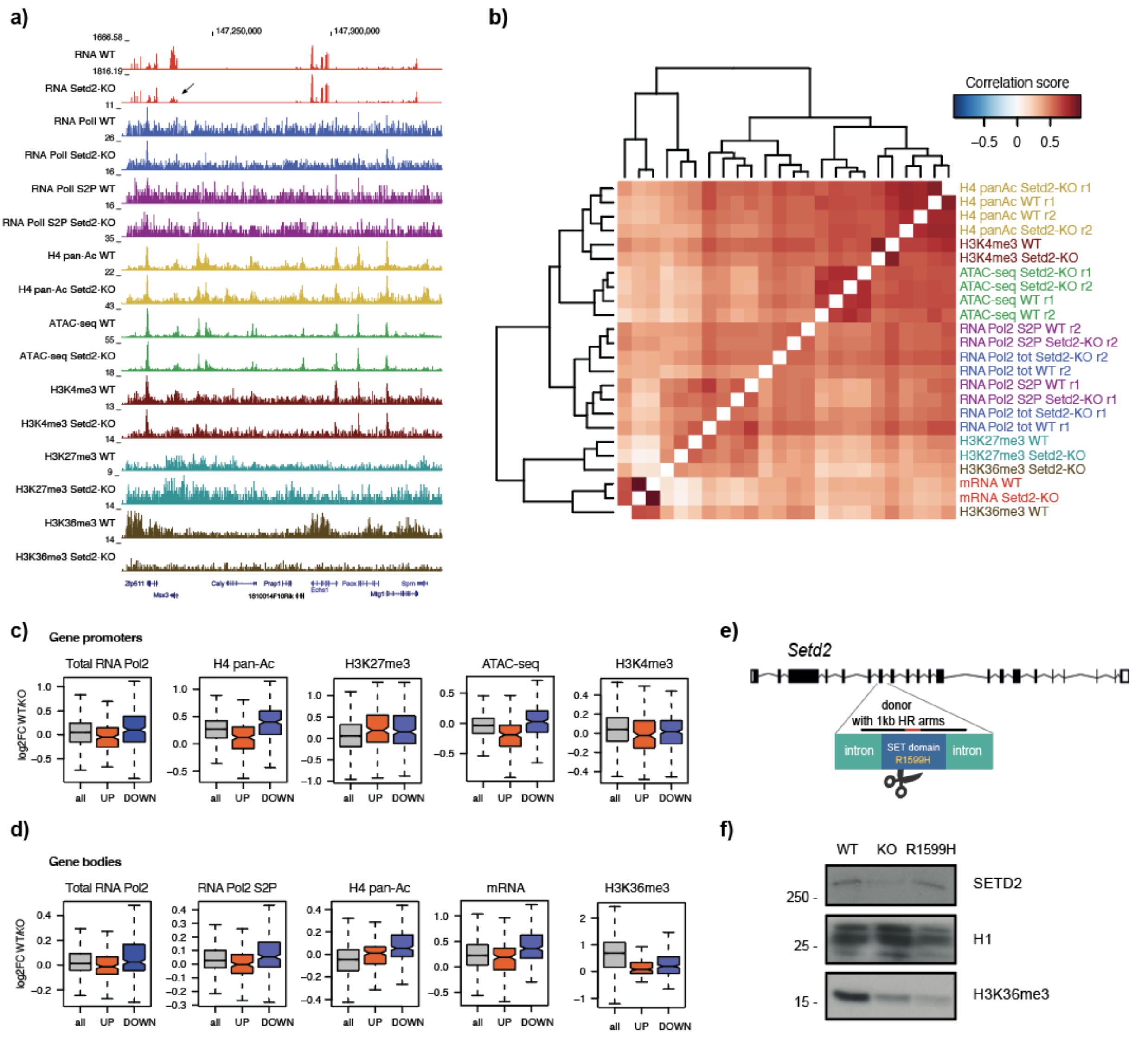


**Figure S6. The catalytic activity of SETD2 is largely dispensable for neuronal differentiation.**

**a)** Representative genome browser view exemplifying ChIP-seq signals of various chromatin marks and ATAC-seq signal between wild-type and *Setd2*-KO NPCs. Shown are read counts per 100 bp for ChIP-seq samples. Arrow indicates downregulation of the neuronal homeodomain transcription factor *Msx3*. **b)** Clustered heatmap presenting correlation scores between various ChIP- and ATAC-seq levels wild-type and *Setd2*-KO NPCs (in replicates). Datapoints are 1kb sized genomic intervals covering the entire genome. **c-d)** Boxplots indicating the differences in various chromatin marks as function of gene expression differences between wild-type and *Setd2*-KO NPCs (c) at promoters and (d) in gene bodies. **e)** Schematic overview for the generation of an mESC line harboring an endogenous mutation at the catalytic SET domain of *Setd2* leading to an amino acid change from arginine (R) to histidine (H) at position 1599. **f)** Western blot indicating a reduction in H3K36me3 in presence of the *Setd2* R1599H mutation. Histone H1 and LAMBIN B1 serve as loading controls.

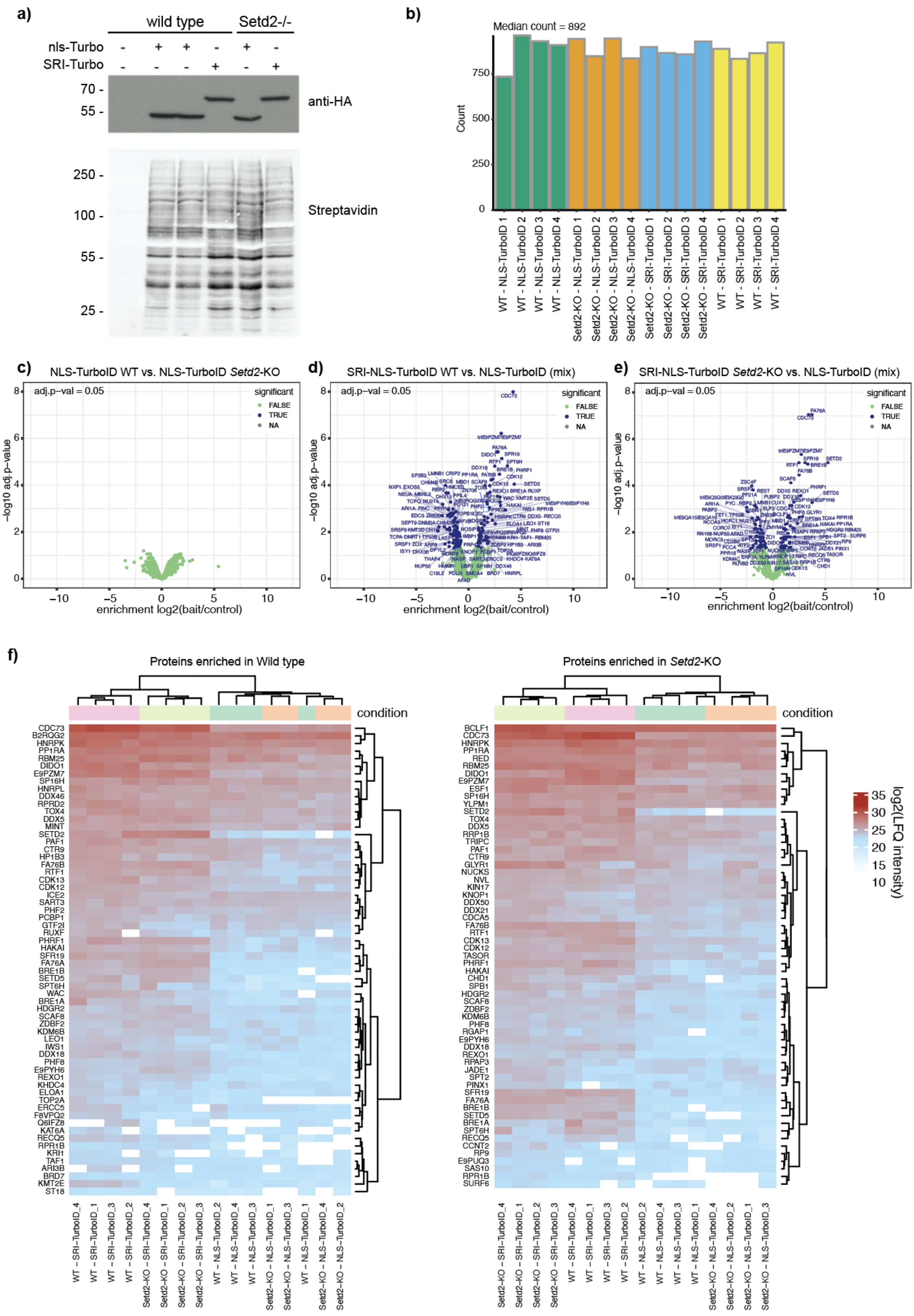
a)
wild type
Setd2-/-
nls-Turbo
SRI-Turbo
70 -
55 -
anti-HA
250 -
100 -
55 -
25 -
Streptavidin
b)
Median count = 892
Count
750
500
250
0
WT - NLS-TurboID 1
WT - NLS-TurboID 2
WT - NLS-TurboID 3
WT - NLS-TurboID 4
Setd2-KO - NLS-TurboID 1
Setd2-KO - NLS-TurboID 2
Setd2-KO - NLS-TurboID 3
Setd2-KO - NLS-TurboID 4
Setd2-KO - SRI-TurboID 1
Setd2-KO - SRI-TurboID 2
Setd2-KO - SRI-TurboID 3
Setd2-KO - SRI-TurboID 4
WT - SRI-TurboID 1
WT - SRI-TurboID 2
WT - SRI-TurboID 3
WT - SRI-TurboID 4
c)
NLS-TurboID WT vs. NLS-TurboID Setd2-KO
d)
SRI-NLS-TurboID WT vs. NLS-TurboID (mix)
e)
SRI-NLS-TurboID Setd2-KO vs. NLS-TurboID (mix)
adj.p−val = 0.05
significant
FALSE
TRUE
NA
−log10 adj.p−value
enrichment log2(bait/control)
f)
Proteins enriched in Wild type
Proteins enriched in Setd2-KO
condition
log2(LFQ intensity)
35
30
25
20
15
10

**Figure S7. SRI-ChromID reveals proteins associated with the elongating RNA Pol II.**

**a)** Western blot analysis shows HA-tagged SRI-TurboID and NLS-TurboID protein expression in wild type and *Setd2*-KO cells (top). Increased protein biotinylation upon presence of TurboID proteins is indicated by Streptavidin detection of biotinylated proteins (bottom). **b)** Number of individual proteins detected in four replicates after proximity biotinylation in the respective cell lines and detection via mass spectrometry. **c)** Volcano plot showing results comparing the nuclear TurboID controls between WT and *Setd2*-KO NPCs. **d-e)** ChromID results showing enriched proteins using SRI-TurboID over the pooled NLS-TurboID samples in wild type (d) and *Setd2*-KO NPC cells (e). Statistically enriched proteins are indicated (FDR-corrected two-tailed *t*-test: FDR = 0.01, s0 = 0.1, $\log_2$ FC > 0, $n = 4$ independent replicates). **f)** Heat map representation of proteins significantly enriched in wild type or *Setd2*-KO NPCs over the pooled NLS-TurboID sample. Shown are the log2-transformed LFQ intensity obtained from all samples and replicates.

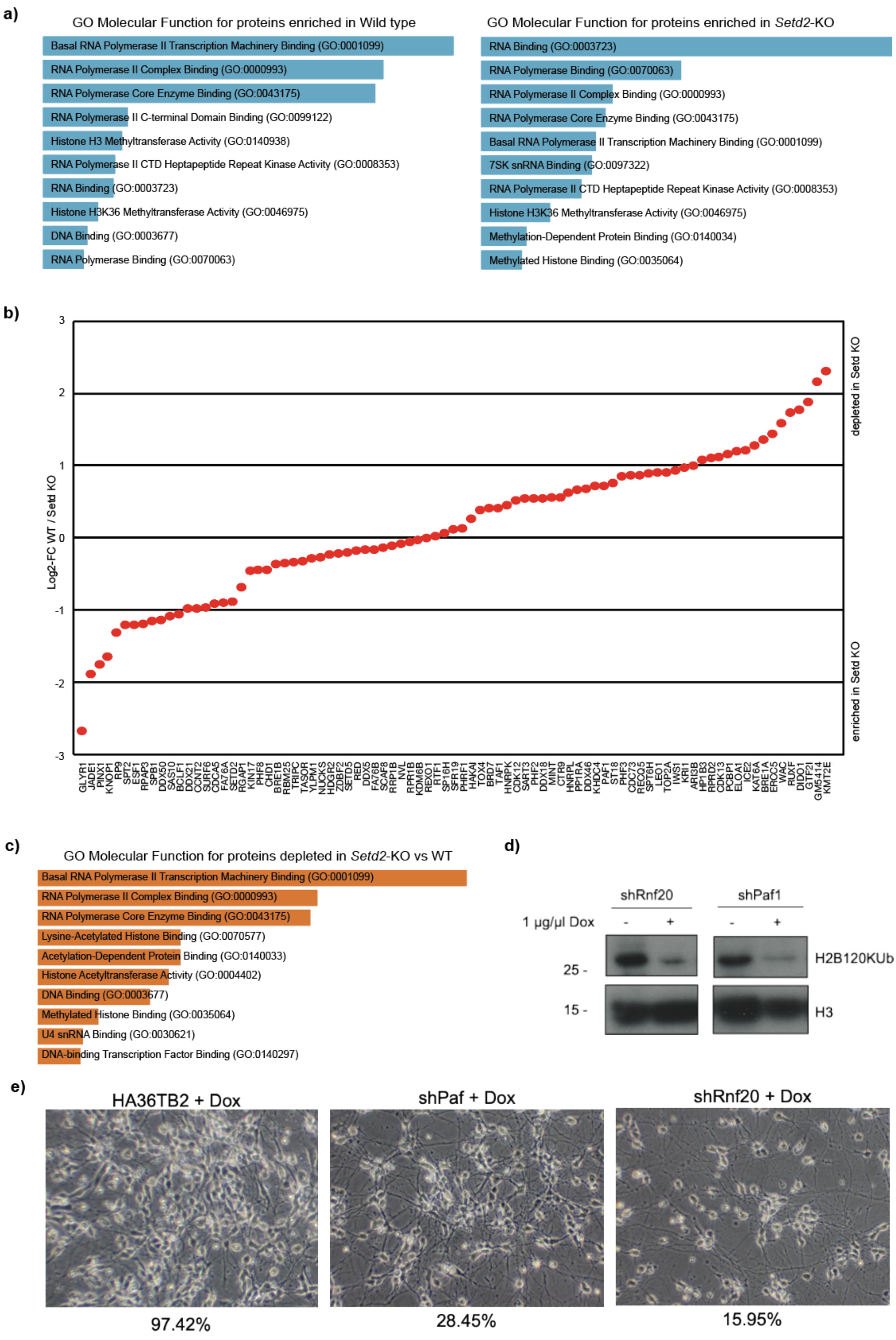
a)
GO Molecular Function for proteins enriched in Wild type
Basal RNA Polymerase II Transcription Machinery Binding (GO:0001099)
RNA Polymerase II Complex Binding (GO:0000993)
RNA Polymerase Core Enzyme Binding (GO:0043175)
RNA Polymerase II C-terminal Domain Binding (GO:0099122)
Histone H3 Methyltransferase Activity (GO:0140938)
RNA Polymerase II CTD Heptapeptide Repeat Kinase Activity (GO:0008353)
RNA Binding (GO:0003723)
Histone H3K36 Methyltransferase Activity (GO:0046975)
DNA Binding (GO:0003677)
RNA Polymerase Binding (GO:0070063)
GO Molecular Function for proteins enriched in Setd2-KO
RNA Binding (GO:0003723)
RNA Polymerase Binding (GO:0070063)
RNA Polymerase II Complex Binding (GO:0000993)
RNA Polymerase Core Enzyme Binding (GO:0043175)
Basal RNA Polymerase II Transcription Machinery Binding (GO:0001099)
7SK snRNA Binding (GO:0097322)
RNA Polymerase II CTD Heptapeptide Repeat Kinase Activity (GO:0008353)
Histone H3K36 Methyltransferase Activity (GO:0046975)
Methylation-Dependent Protein Binding (GO:0140034)
Methylated Histone Binding (GO:0035064)
b)
3
2
1
0
-1
-2
-3
Log2-FC WT / Setd KO
depleted in Setd KO
enriched in Setd KO
GLYR1 JADE1 PINX1 KNOP1 RP9 SPT2 ESF1 RPAP3 SPB1 DDX50 SAS10 BCLF1 DDX21 CCNT2 SURF6 CDCA5 FA76A SETD2 RGAP1 KIN17 PHF8 CHD1 BRE1B RBM25 TRIPC TASOR YLPM1 NUCKS HDGR2 ZDBF2 SETD5 RED DDX5 FA76B SCAF8 RRP1B NVL RPR1B KDM6B REXO1 RTF1 SP16H SFR19 PHRF1 HAKAI TOX4 BRD7 TAF1 HNRPK CDK12 SART3 PHF2 DDX18 MINT CTR9 HNRPL PP1RA DDX46 KHDC4 PAF1 ST18 PHF3 CDC73 RECQ5 SPT6H LEO1 TOP2A IWS1 KRI1 ARI3B HP1B3 RPRD2 CDK13 PCBP1 ELOA1 ICE2 KAT6A BRE1A ERCC5 WAC RUXF DIDO1 GTF2I GM5414 KMT2E
c)
GO Molecular Function for proteins depleted in Setd2-KO vs WT
Basal RNA Polymerase II Transcription Machinery Binding (GO:0001099)
RNA Polymerase II Complex Binding (GO:0000993)
RNA Polymerase Core Enzyme Binding (GO:0043175)
Lysine-Acetylated Histone Binding (GO:0070577)
Acetylation-Dependent Protein Binding (GO:0140033)
Histone Acetyltransferase Activity (GO:0004402)
DNA Binding (GO:0003677)
Methylated Histone Binding (GO:0035064)
U4 snRNA Binding (GO:0030621)
DNA-binding Transcription Factor Binding (GO:0140297)
d)
shRnf20
shPaf1
1 µg/µl Dox
-
+
-
+
25 -
15 -
H2B120KUb
H3
e)
HA36TB2 + Dox
shPaf + Dox
shRnf20 + Dox
97.42%
28.45%
15.95%

**Figure S8. SRI-ChromID reveals proteins associated with the elongating RNA Pol II.**

**a)** Bar plots representing the top ten Molecular Function GO terms summarizing the proteins enriched by the SRI-TurboID in wild type and *Setd2*-KO NPCs. Data was generated using the Enrichr database. **b)** Direct fold change comparison between WT and *Setd2*-KO cells indicates proteins that show enriched or reduced interactions with elongating RNA Pol II in absence of SETD2. Log2-FC was calculated by subtracting the log2FC enrichments in *Setd2*-KO over pooled NLS-TurboID samples from log2FC enrichments in *WT* over pooled NLS-TurboID. Only proteins that were significantly enriched in wild type and/or *Setd2*-KO cells are shown. **c)** Bar plot representing the top ten Molecular Function GO terms summarizing the proteins depleted from RNA Pol II in *Setd2*-KO NPCs. Data was generated using the Enrichr database. **d)** Western blot showing reduced H2BK120ub upon Tet-induced expression of sh RNAs against Rnf20 or Paf1. Histone H3 serves as loading control. **e)** Microscopy images of vitro derived neurons at day d14 showing the effect of Tet-inducible shRNA against *Paf1* or *Rnf20* on neuronal differentiation. Shown are average percentages of survival after dissociation (plated/attached) of three independent experiments; 100x magnification. Cells were treated with 1 µg/ml doxycycline (DOX) from the start of the differentiation.

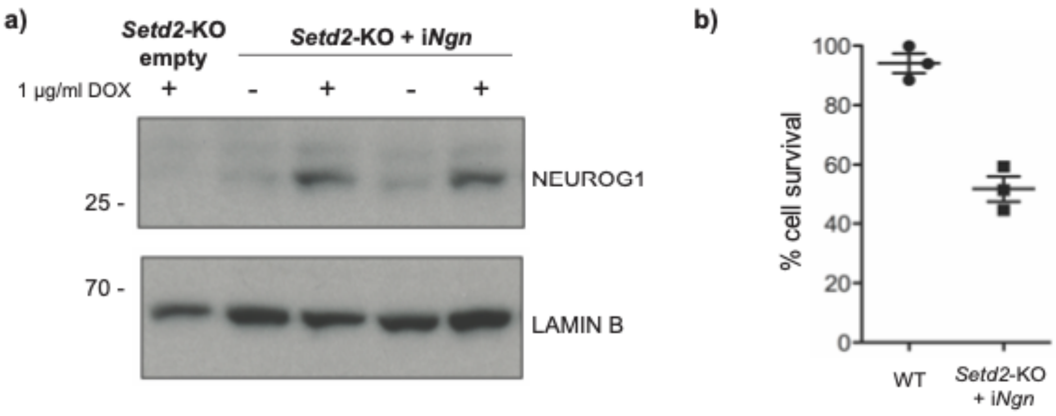


**Figure S9. Overexpression of a neuronal master transcription factors partially rescues cell survival of *Setd2*-deficient NPCs.**

**a)** Immunoblot analysis for Neurogenin 1 (NEUROG1) in two Tet-inducible *neurogenin2-2A-neurogenin1* (i*Ngn*) ESC-derived neurons in comparison to the parental *Setd2*-KO cell line. LAMIN B1 serves as loading control. Cells were treated either without or always with 1 µg/ml doxycycline (DOX) during *in vitro* differentiation. **b)** Shown are percentages of survival at d14 of three independent differentiation rounds.

### Supplementary Tables

**Supplementary Table 1 -** List of differentially expressed genes between Setd2-KO NPC and wild type NPC.

Shown are the DGE signature gene sets in Setd2KO over WT and WT over Setd2KO, respectively. Columns show ENSEMBL, EntrezGene IDs, and Gene symbols. In addition, the logFC, unshrunk logFC, logCPM, PValues and FDR from the edgeR DGE analysis are shown.

**Supplementary Table 2 -** List of significantly identified proteins in ChromID using SRI-BioID in Setd2-KO NPC and wild type NPC.

Shown are the proteins significantly enriched in Setd2-KO and Wild type cells by comparing SRI-TurboID over pooled nTurboID background samples. Columns show protein names, UniprodID, logFC, and qualitative indication if they passed the significance threshold. Statistically enriched proteins are identified by FDR-corrected two-tailed *t*-test: FDR = 0.01, s0 = 0.1, $\log_2$ FC > 0, $n = 4$ independent replicates.

**Supplementary Table 3 -** List of all oligo sequences used in this study.

| | |
|---|---|
| RT-qPCR Sox2 forward | ACAGCATGTCCTACTCGCAG |
| RT-qPCR Sox2 reverse | ATGCTGATCATGTCCCGGAG |
| RT-qPCR Klf4 forward | GCAGTCACAAGTCCCCTCTC |
| RT-qPCR Klf4 reverse | TTTGCCACAGCCTGCATAGT |
| RT-qPCR Nanog forward | GAACTCTCCTCCATTCTGAACCT |
| RT-qPCR Nanog reverse | GACCATTGCTAGTCTTCAACCAC |
| RT-qPCR Oct4 forward | AAGCGAACTAGCATTGAGAACC |
| RT-qPCR Oct4 reverse | CATACTCGAACCACATCCTTCTC |
| RT-qPCR Setd2 forward | TGGGGCCTTCGTGTGCTATG |
| RT-qPCR Setd2 reverse | GCAATCTTCTCCACATGCTACTT |
| RT-qPCR Crabp1 forward | AACTTCAAGGTCGGAGAGGG |
| RT-qPCR Crabp1 reverse | GCTCTCGGGTCCAGTAAGTT |
| RT-qPCR Msx3 forward | GAGTGCGCGACTGGAGG |
| RT-qPCR Msx3 reverse | CACAGAGCACGGACCACTC |
| RT-qPCR Foxd3 forward | GCAACTACTGGACCCTGGAC |
| RT-qPCR Foxd3 reverse | GCTCCGAAGCTCTGCATCAT |
| RT-qPCR Ngn1 forward | AGGAGTCGTCGCGTCAAAG |
| RT-qPCR Ngn1 reverse | TCTTGGTGAGCTTGGTGTCG |
| RT-qPCR Gata4 forward | TGGGGAGATTAGGTGAGGGG |
| RT-qPCR Gata4 reverse | ATTAGCTGCACAACTGGGCT |
| RT-qPCR Fgf4 forward | CTACCGGATAGGAGACCCTTAGA |
| RT-qPCR Fgf4 reverse | CCTCTTGTCCATTCCTAGTTCCT |
| RT-qPCR Sox17 forward | ATGCATTCTGGACCCGCTAC |
| RT-qPCR Sox17 reverse | AGCTCTCCTGCCTCTCAGAA |
| RT-qPCR Pax6 forward | ATCGAAGGGCCAAATGGAGAA |
| RT-qPCR Pax6 reverse | GTGGGATTGGCTGGTAGACA |
| RT-qPCR Pax3 forward | CATGCCCGGGTTCTCTCTTT |
| RT-qPCR Pax3 reverse | GTCCCATGGTTGCGTCTCTA |
| sgRNA Setd2 exon 3 us | TTAGGTCTCTGTAGGAATGGG |
| sgRNA Setd2 exon 3 ds | AGCGTGTCCTCTCACGATAA |
| sgRNA Setd2 SET mutation | AGAGCAATTCCCTTTTTGAG |
| shRNA Setd2 (TRCN0000238536) | ATAGTGTGACCTCGCCTTATT |
